# The disruption of trace element homeostasis due to aneuploidy as a unifying theme in the etiology of cancer

**DOI:** 10.1101/002105

**Authors:** Johannes Engelken, Matthias Altmeyer, Renty B. Franklin

## Abstract

**Abstract for Scientists:** While decades of cancer research have firmly established multiple “hallmarks of cancer” ^1,2^, cancer’s genomic landscape remains to be fully understood. Particularly, the phenomenon of aneuploidy – gains and losses of large genomic regions, i.e. whole chromosomes or chromosome arms – and why most cancer cells are aneuploid remains enigmatic ^3^. Another frequent observation in many different types of cancer is the deregulation of the homeostasis of the trace elements copper, zinc and iron. Concentrations of copper are markedly increased in cancer tissue and the blood plasma of cancer patients, while zinc levels are typically decreased ^4–9^. Here we discuss the hypothesis that the disruption of trace element homeostasis and the phenomenon of aneuploidy might be linked. Our tentative analysis of genomic data from diverse tumor types mainly from The Cancer Genome Atlas (TCGA) project suggests that gains and losses of metal transporter genes occur frequently and correlate well with transporter gene expression levels. Hereby they may confer a cancer-driving selective growth advantage at early and possibly also later stages during cancer development. This idea is consistent with recent observations in yeast, which suggest that through chromosomal gains and losses cells can adapt quickly to new carbon sources ^10^, nutrient starvation ^11^ as well as to copper toxicity ^12^. In human cancer development, candidate driving events may include, among others, the gains of zinc transporter genes *SLC39A1* and *SLC39A4* on chromosome arms 1q and 8q, respectively, and the losses of zinc transporter genes *SLC30A5*, *SLC39A14* and *SLC39A6* on 5q, 8p and 18q. The recurrent gain of 3q might be associated with the iron transporter gene *TFRC* and the loss of 13q with the copper transporter gene *ATP7B*. By altering cellular trace element homeostasis such events might contribute to the initiation of the malignant transformation. Intriguingly, attenuation or overexpression of several of these metal transporter genes has been shown to lead to malignant cellular behavior in vitro. Consistently, it has been shown that zinc affects a number of the observed “hallmarks of cancer” characteristics including DNA repair, inflammation and apoptosis, e.g. through its effects on NF-kappa B signaling. We term this model the “aneuploidy metal transporter cancer” (AMTC) hypothesis and find it compatible with the cancer-promoting role of point mutations and focal copy number alterations in established tumor suppressor genes and oncogenes (e.g. *MYC*, *MYCN*, *TP53*, *PIK3CA*, *BRCA1*, *ERBB2*). We suggest a number of approaches for how this hypothesis could be tested experimentally and briefly touch on possible implications for cancer etiology, metastasis, drug resistance and therapy.

**Abstract for Kids:** We humans are made up of many very small building blocks, which are called cells. These cells can be seen with a microscope and they know how to grow and what to do from the information on the DNA of their chromosomes. Sometimes, if this information is messed up, a cell can go crazy and start to grow without control, even in places of the body where it should not. This process is called cancer, a terrible disease that makes people very sick. Scientists do not understand exactly what causes cells to go crazy, so it would be good to find out. Many years ago, scientists observed that chromosomes in these cancer cells are missing or doubled but could not find an explanation for it. More recently, scientists have detected that precious metals to our bodies, which are not gold and silver, but zinc, iron and copper, are not found in the right amounts in these crazy cancer cells. There seems to be not enough zinc and iron but too much copper, and again, scientists do not really understand why. So there are many unanswered questions about these crazy cancer cells and in this article, we describe a pretty simple idea on how chromosome numbers and the metals might be connected: we think that the missing or doubled chromosomes produce less or more transporters of zinc, iron and copper. As a result, cancer cells end up with little zinc and too much copper and these changes contribute to their out-of-control growth. If this idea were true, many people would be excited about it. But first this idea needs to be investigated more deeply in the laboratory, on the computer and in the hospitals. Therefore, we put it out on the internet so that other people can also think about and work on our idea. Now there are plenty of ways to do exciting experiments and with the results, we will hopefully understand much better why cancer cells go crazy and how doctors could improve their therapies to help patients in the future.

**Abstract for Adults:** One hundred years ago, it was suggested that cancer is a disease of the chromosomes, based on the observations that whole chromosomes or chromosome arms are missing or duplicated in the genomes of cells in a tumor. This phenomenon is called “aneuploidy” and is observed in most types of cancer, including breast, lung, prostate, brain and other cancers. However, it is not clear which genes could be responsible for this observation or if this phenomenon is only a side effect of cancer without importance, so it is important to find out. A second observation from basic research is that concentrations of several micronutrients, especially of the trace elements zinc, copper and iron are changed in tumor cells. In this article, we speculate that aneuploidy is the reason for these changes and that together, these two phenomena are responsible for some of the famous hallmarks or characteristics that are known from cancer cells: fast growth, escape from destruction by the immune system and poor DNA repair. This idea is new and has not been tested yet. We name it the “<u>a</u>neuploidy <u>m</u>etal transporter <u>c</u>ancer” (AMTC) hypothesis. To test our idea we used a wealth of information that was shared by international projects such as the Human Genome Project or the Cancer Genome Atlas Project. Indeed, we find that many zinc, iron and copper transporter genes in the genome are affected by aneuploidy. While a healthy cell has two copies of each gene, some tumor cells have only one or three copies of these genes. Furthermore, the amounts of protein and the activities of these metal transporters seem to correlate with these gene copy numbers, at least we see that the intermediate molecules and protein precursors called messenger RNA correlate well. Hence, we found that the public data is compatible with our suggested link between metal transporters and cancer. Furthermore, we identified hundreds of studies on zinc biology, evolutionary biology, genome and cancer research that also seem compatible. For example, cancer risk increases in the elderly population as well as in obese people, it also increases after certain bacterial or viral infections and through alcohol consumption. Consistent with the AMTC hypothesis and in particular, the idea that external changes in zinc concentrations in an organ or tissue may kick off the earliest steps of tumor development, all of these risk factors have been correlated with changes in zinc or other trace elements. However, since additional experiments to test the AMTC hypothesis have not yet been performed, direct evidence for our hypothesis is still missing. We hope, however, that our idea will promote further research with the goal to better understand cancer – as a first step towards its prevention and the development of improved anti-cancer therapies in the future.

## Introduction

### Genomic landscape of cancer

Despite the widespread view from the clinics that "cancer is not just one disease but many diseases" (National Cancer Institute at NIH; http://www.cancer.gov), there exist striking similarities between different types of cancer. On the functional level, a number of shared hallmarks across different types of cancer have been described ^1,2^. On the genomic level, certain chromosomal gains and losses cluster non-randomly to the same genomic regions and a number of shared, recurrently mutated genes and pathways point to significant similarities between different types of tumors and the way they develop. Interestingly, the opposite can also be observed, namely tumors with very different combinations of chromosomal gains and losses or point mutations that still converge on the same hallmark phenotype. This raises the important question whether there exists a (currently missing) unifying theme or underlying molecular mechanism for carcinogenesis across all types of cancer.

Overall, the genomic landscape of cancer is highly complex as reviewed elsewhere ^13^ and includes different types of point mutations, focal amplifications and deletions as well as non-exclusive chromosome-level phenomena such as aneuploidy ^3^, whole genome duplications leading to a tetraploid genome and subsequent losses ^14,15^, translocations such as the Philadelphia chromosome ^16^, and massive rearrangements such as the observed chromosome shattering in chromothripsis ^17^. Recently a subdivision of tumor types into two groups dominated by either point mutations or copy number variants was suggested ^18^.

Aneuploidy is a frequent observation across virtually all types of cancer and a characteristic of all solid tumors (e.g. Supplemental Figure 7 in ^19^, Figure 3 in ^20^ and Supplemental Figure S1 (Supplemental Note A). The works of David Paul von Hansemann ^21^ that were later taken up by Theodor Heinrich Boveri ^3^ recognized many years ago that an incorrect number of chromosomes was caused by errors during mitosis and that this was related to the cancerous state of a cell ^22^. Boveri already saw cancer as a disease of the chromosomes ^3,23^ and suggested that aneuploidy was causal for cancer development. Before, Walter Sutton had proposed the chromosome theory of inheritance ^24^. The idea of a causal role of aneuploidy was later restated several times, most prominently in recent years by Peter Duesberg ^25,26^ and it was suggested that aneuploidy could, by deregulating large numbers of genes, entail the first predispositions for eventual cancer development. However, whether aneuploidy is indeed causally related to cancer development has remained controversial and is still one of the big open questions in cancer biology.

Interestingly, general support for the idea of aneuploidy as a means for cell adaptation can be found in studies on yeast. Large-scale chromosomal rearrangements seem indeed capable of driving quick adaptation of trace element homeostasis, as was first observed in the adaptive response to a new carbon source ^10^. More recently, this was demonstrated in yeast for copper tolerance; specifically, chromosomal aberrations including the transcriptional activator CUP2 and the metallothionein CUP1 seemed to drive this phenomenon ^12^. Similar genomic rearrangements were able to drive resistance to nutrient starvation in yeast ^11,27^. Translated to the process of clonal evolution of cancer cells, these examples show that chromosomal gains and losses may indeed quickly lead to changes in trace element homeostasis. Comparisons between aneuploidy in yeast and cancer tumors are consistent with the idea of aneuploidy as a driving force of adaptation and also leave space for the existence of a selective force ^28,29^.

Regarding gene copy number variation, in this article we consider chromosome arm-level and focal events as separate, independent phenomena. This is consistent with a recent study ^19^ which found that across 17 different tumor types, approximately 35% of the genome was affected by any aberration: while 25% of the genome was affected by chromosome arm aberrations, only 10% was affected by focal aberrations – importantly with no more than 2% overlap. Moreover, this discrimination is also compatible with the observation that background mutation rates for arm-level and focal events are significantly different ^30^.

### Trace elements in cancer

It is well established that the homeostasis of trace elements such as zinc, iron and copper is disrupted in diverse cancers both on the intracellular and on the tissue level. This is most clearly demonstrated by the fact that, in different epithelial cancers, tissue zinc concentrations are significantly reduced when compared to their healthy tissue counterparts. For example, zinc is reduced in malignant liver tissue ^6,9^, zinc is low and copper is high in malignant lung tissue ^5^ and in cervical cancer tissue ^31^, while in breast cancer, tissue zinc levels are strongly increased^4^. Garg et al. show the deregulation of many trace elements in breast cancer tissue ^32^. Copper is markedly increased in glioma but not in the less dangerous meningioma ^33^. Gupte and Mumper ^34^ present an overview on copper in serum and diverse malignant tissues. Additional links between trace element homeostasis and cancer exist (Supplemental Note C).

Whether these changes in tissue trace element concentrations are causal for malignancy or whether they are a consequence of other cellular changes that occur during carcinogenesis is an important unresolved question. For example, zinc deficiency has been suggested to cause oxidative DNA damage, to disrupt p53, NFkB, and AP1 DNA binding and hereby compromise the cellular capability to deal with stress ^35^. Other authors ^7^ would argue that such changes in zinc may provide an intracellular environment for the expansion of transformed cells. In support of this view, it was observed that zinc levels change gradually with the grade of prostate cancer ^36^. Interestingly, low zinc concentrations likely precede malignancy in prostate gland cells ^37^. This last point supports the idea that changes in zinc homeostasis may indeed be causal contributing factors rather than consequences in the progression towards cancer.

### Objectives of this article

The main objectives of this hypothesis article are to (i) explain the AMTC hypothesis in a framework of clonal cancer evolution, (ii) discuss its compatibility with mRNA expression and DNA copy number variation data derived from diverse cancer types, and (iii) discuss a possible role of trace element homeostasis in the established framework of cancer hallmarks. Further, (iv) we provide a short overview how various aspects of the AMTC hypothesis could be tested in different experimental model systems (Text Box B). Finally, (v) we address possible implications for cancer risk factors, metastasis and drug resistance (Appendix).

## Model of carcinogenesis

### The aneuploidy metal transporter cancer (AMTC) hypothesis

As our main hypothesis, we suggest that expression changes of genes encoding different metal transporters may underlie, i.e. drive the selection of recurrently observed chromosomal gains and losses in cancer. Such gene dosage effects may disrupt the cellular trace element homeostasis and thereby contribute to cancer development and even affect the established hallmarks of cancer (Figure 1). This would unveil a completely novel and unexplored link between aneuploidy and the observed disruption of trace element homeostasis in cancer, and therefore we term it the “aneuploidy metal transporter cancer” (AMTC) hypothesis. Limitations and complexities of this hypothesis are discussed in Supplemental Note B.

**Figure 1:**
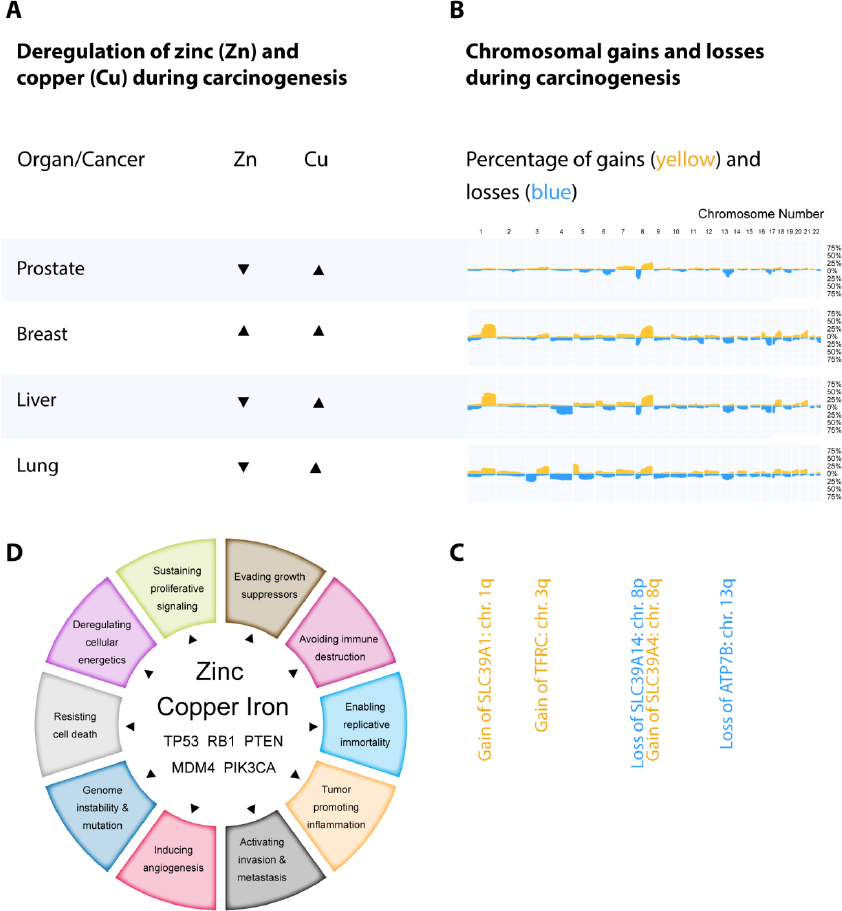
<u>A</u>neuploidy <u>M</u>etal <u>T</u>ransporter <u>C</u>ancer (AMTC) Hypothesis. (A) Trace element concentrations are deregulated in different tumors, which typically show lower zinc and higher copper when compared to normal tissue. (B) Recurrent chromosomal gains leading to tumor samples with three copies of a particular chromosome arm (orange) or losses (one copy, blue); percentages of gains and losses across many samples are visualized along the genome (http://progenetix.org). Based on (A) previous evidence of deregulated trace element concentrations and (B) characteristic chromosomal gains and losses (aneuploidy) in diverse types of cancer, we hypothesize that there may be a causal link between these two observations, i.e. (C) gains and losses of specific chromosome arms that include metal transporters may lead to the disruption of trace element homeostasis during carcinogenesis, which in turn might contribute to cancer progression. (D) The disruption of trace element homeostasis likely affects several of the established hallmarks of cancer and hereby, together with established recurrently mutated cancer genes, may contribute to malignant transformation. If this hypothesis were to be confirmed, the disruption of trace element homeostasis would emerge as a unifying theme in cancer research.

If the AMTC hypothesis were correct and metal transporter gene dosage effects would underlie chromosomal gains and losses in cancer, the disruption of intracellular trace element homeostasis should roughly coincide with the emergence of aneuploidy. Indeed, both phenomena seem to occur early in the course of carcinogenesis. For example, changes in zinc concentrations were found as early events in three studies of benign prostatic hyperplasia ^7^, although more research on trace element concentrations in pre-cancerous lesions is clearly needed. Aneuploidic changes have likewise emerged as an early event in carcinogenesis across many types of cancer ^38–41^. In fact, chromosomal aberrations are not only common in malign and metastatic tumors but already in early lesions and benign tumors ^42–49^. Cytogenetic methods, which were greatly improved by the human genome project ^50,51^, made these studies possible. They are in agreement with a recent study based on resequencing technology which found that early neoplasiae in breast tissues often show aneuploidy of characteristic chromosome arms but only show very few point mutations ^52^.

Is there a functional basis for the AMTC hypothesis? Costello and Franklin ^53,54^ have alluded to a possible causal role of zinc in prostate cancer. A role for zinc homeostasis in the etiology of breast cancer has also been proposed ^55^. Importantly, as we discuss below, zinc may play a role in diverse hallmark characteristics of cancer. Further, we note that a number of experimental studies on specific metal transporters seem compatible with their suggested role as driver genes. For example, the overexpression of zinc transporter ZIP4 (SLC39A4) increased cell proliferation in vitro and enhanced tumor growth in a mouse xenograft model ^56^ whereas attenuation of ZIP4 led to a number of cancer inhibiting effects such as decreased cell migration and enhanced apoptosis in vitro and slower tumor growth in a mouse xenograft model ^57,58^. The iron transport protein TFRC (transferrin receptor protein 1) showed an interaction with c-Myc and its overexpression lead to increased proliferation ^59^, while suppression of TFRC by antisense RNA lead to the inhibition of tumor growth ^60^. Interestingly, cells overexpressing the copper transporter ATP7B outcompeted untransduced control cells in high copper cell culture conditions ^61^, but it seems unclear whether this could indicate malignant behavior. In partial agreement with the AMTC hypothesis, suppression of the zinc transporter ZIP6 (SLC39A6, LIV-1) inhibits HeLa cell invasiveness ^62^, whereas attenuation of ZIP6 promotes the epithelial-to-mesenchymal transition in ductal breast tumor cells ^63^. Interestingly, overexpression of the zinc transporter ZIP1 (SLC39A1) inhibits tumor growth in prostate cancer cells ^64,65^. Of note, prostate cancer cells typically do not show a gain of chromosome arm 1q, which encodes SLC39A1. In summary, most of the observed effects correlate well with the direction of change that would be expected from the observed recurrent gains and losses of the corresponding chromosome arms (i.e. *SLC39A4* +8q;*TFRC* +3q, *SLC39A6* +18q).

### Evolutionary dynamics of early carcinogenesis

While recent achievements in whole genome sequencing have defined the scope and quality of mutations and genomic rearrangements in various advanced human tumors, our understanding of tumor evolution and how early genomic changes contribute to the initial clonal expansions that can then result in tumor development remains in its infancy. Several excellent reviews on cancer genome evolution ^66–70^ have emphasized the evolutionary aspects of cancer development. Here, we have made an attempt to understand the trace element-centric view provided by the AMTC hypothesis in the context of clonal tumor evolution (Figure 2).

**Figure 2:**
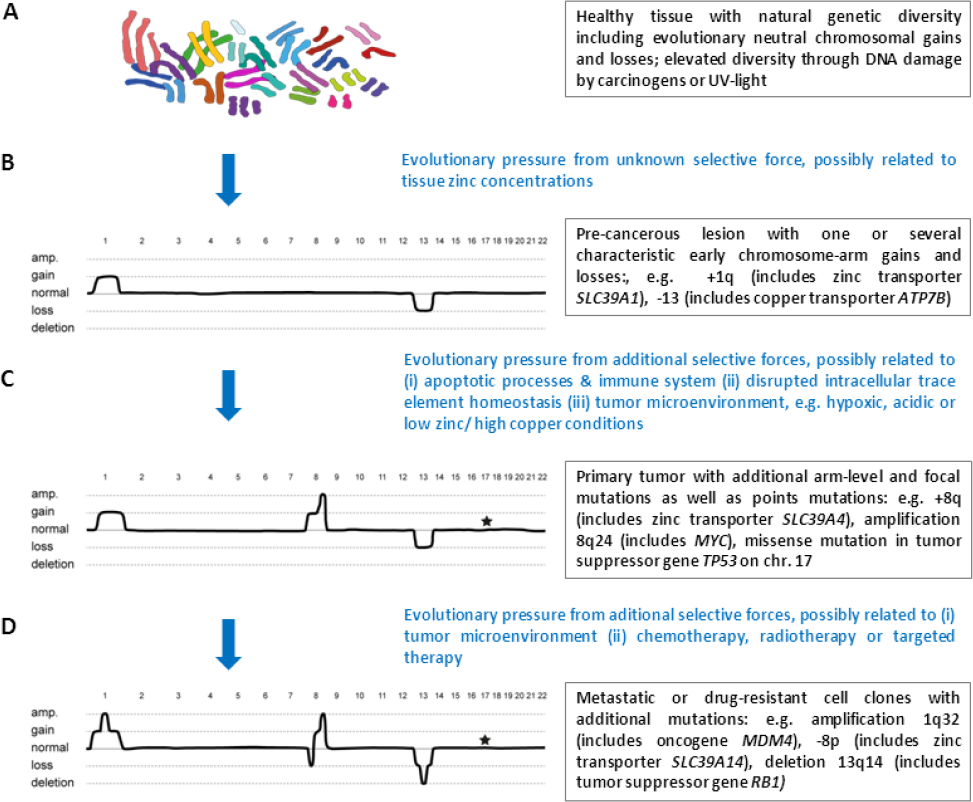
Tentative evolutionary view of carcinogenesis, exemplified by a single evolving cancer genome. (A) Healthy tissue contains random, somatic chromosome-arm mutations in a small subset of cells. (B) Evolutionary pressure, possibly positive selection for cell clones able to grow in tissue affected by localized zinc deficiency (which occurs e.g. during certain bacterial infections), favors clones with specific chromosome gains (three copies) or losses (one copy). Similar adaptive chromosomal mutations have been observed in populations of yeast cells. As the precancerous cell clone grows, it becomes increasingly transformed by acquiring certain hallmark characteristics. As discussed in the main text, these hallmarks could be intertwined with intracellular changes in trace element homeostasis. (C) Point mutations as well as focal amplifications (multiple copies) or deletions (zero copies) of numerous established cancer genes may provide a selective growth advantage during later stages of carcinogenesis. (D) Subsequent events, possibly affecting additional metal transporters, may underlie the progression to metastasis. Hereditary cancers may include similar mutations and selective pressures, but in a somewhat different order.

From an evolutionary point of view, it seems justified to assume that cancer never starts from a mutation alone, but always from a combination of a pre-disposing mutation in combination with an early selective force. This is consistent with the observation that exposure to a DNA damaging carcinogenic substance may not be enough to cause cancer ^71^. In the context of the AMTC hypothesis, we assume that the gain or loss of metal transporters is not sufficient to cause cancer but a selective force is needed to start the evolutionary process and escape normal growth control. In fact, it seems quite possible that potentially carcinogenic cells may remain in a dormant state until the microenvironment in a tissue changes in such a way that these cells acquire an advantage in fitness and start to clonally expand. It is tempting to speculate that the putative selective force is related to trace element homeostasis itself (Figure 2).

Hence, might zinc deficiency or some other deregulated trace element in the surrounding tissue or blood stream act as an early selective force? Several observations fit into this picture: zinc deficiency promotes esophageal cancer after carcinogen induction ^72^ and replenished zinc protects against this effect ^73,74^. This also fits with the observation that zinc deficiency induced inflammation contributes to the development of esophageal cancer in rats and that carcinogen-exposure without zinc deficiency was not enough to produce cancer ^71^. Further, zinc is protective against induced carcinogenesis in colon ^75^. The localized area in the tissue where our proposed selective force acts may roughly correspond to the pre-cancerous field that was proposed 60 years ago ^76,77^.

Processes of natural selection ^78,79^ depend on the Malthusian principle of population growth ^80^. In clonal tumor evolution, this prerequisite is possibly represented by cancer stem cells, which have the potential for unlimited cell divisions. Combined with limited resources, this leads to competition between cells. However, healthy cells in a tissue do not usually compete with each other in a Darwinian fashion as opposed to cells in a clonal yeast population or individuals in plant and animal populations. Instead, the number and localization of cells in a tissue is controlled through proliferative and anti-proliferative processes. While strongly mutated or aberrantly behaving cells usually undergo cell death and are thereby eliminated from the population, pre-cancerous cells somehow must overcome this control mechanism before they can compete with each other or out-compete normal cells. Further, phenotypic diversity as a substrate for a selective force to act upon is required. In this evolutionary game, one may expect that the risk that a cancer cell clone emerges is a function of the load of random mutations in a tissue multiplied by the exposure to selective pressures with certain selection coefficients and exposure times. Carcinogens as well as random errors during DNA replication and mitosis lead to the accumulation of mutations in a healthy tissue, while purifying selection can be expected to act against such mutations. The reason for purifying selection is that chromosomal duplications and other mutations are likely to carry a fitness cost in normal tissue, as was observed in cadmium-resistant yeast strains when exposed to low-cadmium environments ^81^. Fitting with the idea of a fitness cost, chromosomal duplications were found as an adaptive but transient response of yeast to different environmental stresses and when the selective pressure was reduced, the adaptive processes were driven by mutations other than chromosomal duplications ^82^. Depending on additional factors such as genetic drift, genetic predisposition and generation time of cells in a tissue, such processes may or may not lead to cancer in the shorter or longer run.

The process of malignant transformation as shown in Figure 2 entails numerous point and focal mutations in known cancer genes. Given the premise that the disruption of trace element levels is a relatively early event, one may also ask whether some of these mutations might be a response to this previous event. As discussed below, especially intracellular zinc levels may influence a number of these pathways. Certainly, the tumor microenvironment is highly complex and numerous known and unknown selective forces beyond trace elements may shape the adaptive response of a tumor cell. The progression of a tumor towards metastasis again may likewise entail several different selective forces and mutations such as chromosome arm gains and losses possibly including metal transporters as well as a number of established recurrent other mutations (Figure 2). Speculatively, some of these mutations may be compensatory mutations ^83^ that keep the cell alive but which compensate the effect of some earlier mutations possibly related to trace element deregulations. Of note, six out of eleven recently confirmed high-confidence cancer driver genes ^84^ are either metal-binding proteins and/or are regulated by metal ions, and their cancer-related function has been linked to trace element homeostasis before: *TP53* ^85–91^, *KEAP1* ^92,93^, *NFE2L2* (also called *NRF2*) ^94,95^, *PTEN* ^96^, *MLL2* ^97^ and *NOTCH1* ^98^. In other words, it seems possible that the first aneuploidic changes might occur even before certain point or focal mutations in established cancer genes arise, the latter potentially representing compensatory responses.

Importantly, this model (Figure 2) may help explain key steps in the progression of hereditary and non-hereditary cancer types alike, which occur in very different incidence rates ^99^ and which both show aneuploidy: While it seems likely that the suggested chromosomal gains and losses including metal transporters would also form part of the carcinogenesis process in hereditary cancers, a mutation in a tumor suppressor gene such as *TP53*, *BRCA1* or *PTEN* is encoded in the somatic genome already and does not need to evolve *in situ*. Therefore, the long process of carcinogenesis may be accelerated in hereditary cancers.

### Metal transporters as strong candidate driver genes

Large scale efforts such as the International Cancer Genome Consortium and the Cancer Genome Atlas including the Pan-Cancer project ^100^ are facilitating the comparative analysis of multi-dimensional datasets across different cancer types. Moreover, thousands of comparative genomic hybridization (CGH) studies have accumulated a rich body of data regarding chromosomal aberrations across many more different cancers and tens of thousands of specimen and are accessible through databases of cytogenetic changes ^101–104^. However, the identification of underlying driver genes through functional studies or fine-mapping efforts has so far proven difficult ^52^. While the causal influence of a number of recurrent events such as point mutations in *TP53* or the high-level amplifications of certain regions including *MYC* or *ERBB2* have been firmly established, it has remained challenging to establish the identity or even existence of driver genes behind recurrent chromosome-arm level gains and losses.

We have carried out a tentative analysis of integrative genomic datasets available at cBioPortal for Cancer Genomics ^105,106^. Methodological details are given in Supplemental Note A. Specifically, we have analyzed data on copy number and mRNA expression from fifteen published datasets including twelve different epithelial carcinomas. We have analyzed 458 genes in different groups: (i) zinc, iron and copper transporter and related genes; (ii) SLC transporter genes as controls (N=326); and (iii) a group of established cancer genes (N=58) mainly from the Cancer Gene Census ^107^, that were enriched in large deletions or in amplifications. For each gene and cancer type, we have extracted three pieces of information: (i) maximum percentage of hemizygous gains or losses; (ii) the absolute log ratio of gains/losses; and (iii) gene dosage. The analysis of gene dosage is visualized in Figure 4 and Supplemental Figures 2A and 2B. The rationale behind these three pieces of information is as follows: (i) The observed frequency of heterozygous gains or losses measures how often a particular chromosomal event recurs in independent cancer specimen and therefore is a key parameter; amplifications (multiple gene copies) and deletions (zero gene copies) may represent more local events and do not strongly overlap with chromosome-arm events ^19^ and therefore were not considered here; (ii) Characteristically, in a certain cancer type, a chromosome arm is either hemizygously lost or gained but does not show both types of aberrations at high frequency; the log ratio of the two frequencies simply measures this characteristic; (iii) Following the assumption that chromosomal gains and losses act mainly by increasing or decreasing the mRNA expression level of one or several encoded genes, we use gene dosage as a criterion to include or exclude genes as potential driver genes in a certain type of cancer. Different statistics and “aneuploidy scores” were calculated for all genes across all epithelial cancers based on these three pieces of information as described in Supplemental Note A.

Strong candidate loci that may drive chromosomal gains and losses in epithelial cancers are listed in Table 1A (full results in Supplemental Table S2). Strikingly, certain metal transporters (e.g. *SLC39A1*, *SLC39A3*, *SLC39A4*, *SLC39A6*, *SLC39A14*, *SLC30A5*, *ATP7B*, *TFRC* among others) show higher aneuploidy scores than many of the cancer census genes. Some metal transporters show only high scores for a subset of cancer types, e.g. *SLC39A3* in uterine, ovarian and lung cancer or *SLC39A7* and *TFR2* in melanoma (score_1_corrected in Supplemental Table S3). Of note, a number of cancer census genes also show high scores (Table 1B), which possibly reflects that their functional effects are indeed related to gene dosage. However, interestingly, for many of these genes, the underlying type of mutation is firmly established, namely through point mutations, focal amplification or deletion rather than chromosome-arm level aberrations. Certainly this leaves the possibility open that some of the established cancer census genes may also act through arm-level mutations. Further reinforcing the differences with the metal transporters, many cancer census genes show an enrichment of point mutations as detected by intOgen ^108^. Once the results are ordered by chromosome arm (Supplemental Table S3), this broad picture is confirmed: the high-ranking candidate genes for most chromosome arms are either (i) established tumor suppressor genes or oncogenes typically associated with focal events or point mutations, or (ii) metal transporters. In summary, this suggests that certain metal transporters may act as drivers in carcinogenesis by gene dosage through gains and losses on the chromosome-arm level, while the established cancer genes mainly act through their sharply defined point and focal mutations.

**Table 1A.** Metal transporters as strong candidate driver genes underlying aneuploidy. In summary, we propose that a number of metal transporter genes, especially ATP7B, TFRC, SLC39A1, SLC39A4, SLC39A14, SLC39A3 and SLC39A6 are candidate driver genes associated with aneuploidy and carcinogenesis. An aneuploidy score based on the gain/loss patterns and gene dosage effects in twelve different epithelial cancer types mainly from the Cancer Atlas Project was calculated for each gene. Complete results are given as Supplemental Data. The column “dosage effects …” refers to the number of datasets in which a particular gene showed a dosage effect in mRNA expression and “tendency of gains or losses” is the predominantly observed gain or loss pattern. For several top candidate driver genes, experimental *in vivo* or *in vitro* overexpression/ downregulation studies have shown malignant or benign phenotypic changes (see main text); the phenotype was either compatible with the predominant chromosomal gain or loss in the candidate gene (“supportive”), “partially supportive” or alternatively designated as “conflictive”. “Aneuploidy driving potential” indicates the manual assignment as likely driver or passenger genes, dependent on the aneuploidy scores in relation to the control group of SLC transporters and to other candidate genes on the same chromosome arm.

| symbol | group | chr | cytoband | focal SCNAs | point mutations | tendency of gains or losses | dosage effects out of 12 epithelial cancers | aneuploidy score_2 across epithelial cancers | experimental | aneuploidy driving potential across epithelial cancers |
| --- | --- | --- | --- | --- | --- | --- | --- | --- | --- | --- |
| - | SLC control mean | - | - | - | - | - | - | 1.643 |  |  |
| ATP7B | copper transport | 13 | q14.3 | - | - | loss | 7 | 3.486 |  | high |
| SLC31A2 | copper transport | 9 | q32 | - | - | loss | 8 | 2.636 |  | med |
| SLC31A1 | copper transport | 9 | q32 | - | - | ambiguous | 9 | 0.832 |  | low |
| ATP7A | copper transport | X | q21.1 | - | point | ambiguous | 1 | 0.216 |  |  |
| TFRC | iron transport | 3 | q29 | - | - | gain | 7 | 5.822 | supportive | high |
| TFR2 | iron transport | 7 | q22.1 | - | - | gain | 2 | 1.146 |  | med |
| SLC11A2 | iron transport | 12 | q13.12 | - | - | ambiguous | 5 | 0.784 |  | low |
| SLC40A1 | iron transport | 2 | q32.2 | - | - | gain | 1 | 0.408 |  | low |
| SLC11A1 | iron transport | 2 | q35 | - | - | ambiguous | 0 | NA |  | low |
| SLC30A5 | zinc efflux ZnT | 5 | q13.1 | - | - | loss | 12 | 4.736 |  | high |
| SLC30A10 | zinc efflux ZnT | 1 | q41 | - | - | gain | 2 | 4.418 |  | low |
| SLC30A1 | zinc efflux ZnT | 1 | q32.3 | - | - | gain | 5 | 3.557 |  | low |
| SLC30A9 | zinc efflux ZnT | 4 | p13 | - | - | loss | 11 | 2.856 |  | med |
| SLC30A6 | zinc efflux ZnT | 2 | p22.3 | - | - | gain | 10 | 2.068 |  | low |
| SLC30A3 | zinc efflux ZnT | 2 | p23.3 | - | - | gain | 6 | 1.850 |  | low |
| SLC30A4 | zinc efflux ZnT | 15 | q21.1 | - | - | loss | 5 | 1.812 |  | low |
| SLC30A7 | zinc efflux ZnT | 1 | p21.2 | - | - | loss | 8 | 1.753 |  | low |
| SLC30A2 | zinc efflux ZnT | 1 | p36.11 | - | - | loss | 1 | 1.244 |  | low |
| SLC30A8 | zinc efflux ZnT | 8 | q24.11 | - | - | ambiguous | 0 | NA |  | low |
| SLC39A1 | zinc influx ZIP | 1 | q21.3 | - | - | gain | 10 | 8.882 | conflictive | high |
| SLC39A4 | zinc influx ZIP | 8 | q24.3 | - | - | gain | 9 | 7.447 | supportive | high |
| SLC39A14 | zinc influx ZIP | 8 | p21.3 | - | - | loss | 9 | 4.797 |  | high |
| SLC39A3 | zinc influx ZIP | 19 | p13.3 | - | - | loss | 8 | 3.150 |  | high |
| SLC39A6 | zinc influx ZIP | 18 | q12.2 | - | - | loss | 10 | 2.748 | part. supportive | high |
| SLC39A8 | zinc influx ZIP | 4 | q24 | - | - | loss | 6 | 2.139 |  | low |
| SLC39A9 | zinc influx ZIP | 14 | q24.1 | - | - | loss | 9 | 1.861 |  | low |
| SLC39A5 | zinc influx ZIP | 12 | q13.3 | - | - | gain | 2 | 1.290 |  | low |
| SLC39A11 | zinc influx ZIP | 17 | q25.1 | - | - | gain | 9 | 1.250 |  | low |
| SLC39A7 | zinc influx ZIP | 6 | p21.32 | - | - | ambiguous | 9 | 1.090 |  | low |
| SLC39A10 | zinc influx ZIP | 2 | q32.3 | - | - | ambiguous | 8 | 0.533 |  | low |
| SLC39A13 | zinc influx ZIP | 11 | p11.2 | - | - | ambiguous | 8 | 0.453 |  | low |
| SLC39A12 | zinc influx ZIP | 10 | p12.33 | - | - | loss | 2 | 0.437 |  | low |
| SLC39A2 | zinc influx ZIP | 14 | q11.2 | - | - | ambiguous | 2 | 0.295 |  | low |

**Table 1B.**
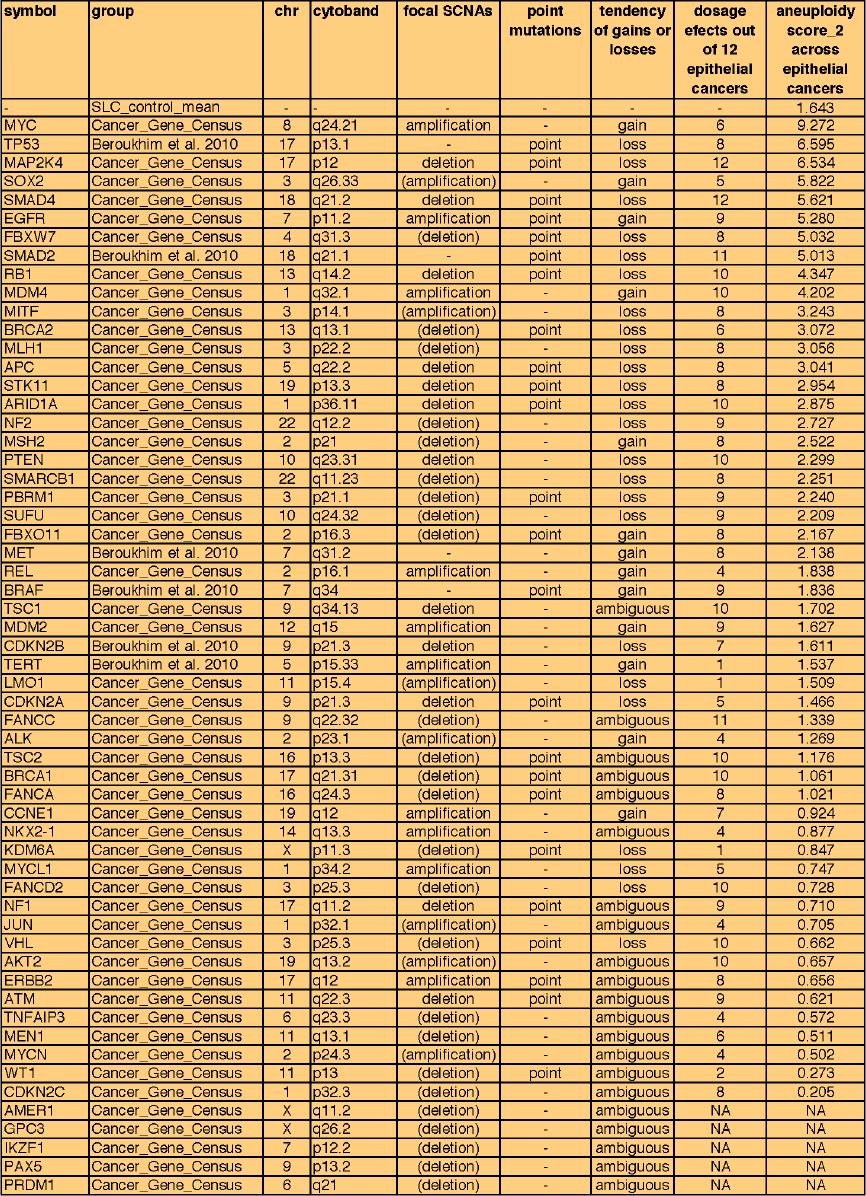
Established cancer genes for comparison. Aneuploidy scores were calculated as for Table 1A. “point_mutations” indicates significant enrichment for point mutations identified either by oncodriveFM, oncodriveCLUST, MutSig or MuSiC. “Focal SCNAs” (somatic copy number alterations) indicates whether significant recurrent focal amplifications or deletions were observed, either in the Cancer Gene Census (shown in brackets) or in the TCGA pan-cancer analysis across 12 solid tumors (shown without brackets). As opposed to the metal transporter genes, many of the established cancer genes show high level amplifications or deletions (the reason for which they were included) as well as an excess of point mutations.

Next, the two families of zinc transporters, ZIP encoded by the *SLC39A* gene family and ZnT encoded by the *SLC30A* gene family, and the control group of the remaining SLC transporters and the Cancer Census Genes were compared to each other across different epithelial cancer types mainly from the Cancer Genome Atlas Project (Figure 3). On the group level, the cancer census genes as well as the two groups of zinc transporters show higher scores than the control group of SLC transporters. Especially, the zinc influx transporters (ZIP) stand out with the highest median value. This effect seems to be caused by relatively few high-scoring genes. On the functional level, the group of ZIP transporters is important in the transport of zinc (and iron) across the cell membrane in epithelial tissues, which may possibly explain why they turned out as strong candidate genes underlying the chromosomal gains and losses in carcinomas. In non-epithelial malignancies different metal transporters or driver genes might be more important (Supplemental Table S3). It remains to be seen whether this or a similar hypothesis can be developed for certain types of leukemia and lymphoma, which typically show absence or low levels of aneuploidy.

**Figure 3:**
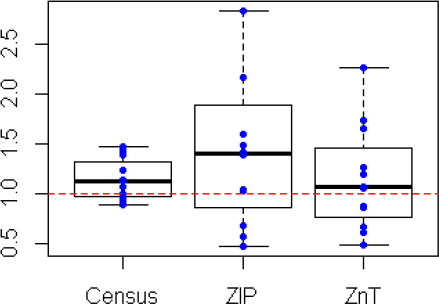
Comparison of different groups of candidate genes underlying aneuploidy. From 12 published datasets of epithelial types of cancer, mean aneuploidy scores were plotted for three groups: a subset of established cancer genes with amplifications or deletions mainly from the Cancer Gene <u>Census</u> ^107^ (median score=1.13, N=58), the <u>ZIP</u>/SLC39A zinc influx transporter family (median score=1.41, N=14 genes) and the <u>ZnT</u>/SLC30A zinc efflux transporters (median score=1.06, N=10). The <u>control group</u> of SLC transporters (median scores set to 1.0, N=326) is displayed as red line. The relatively high mean aneuploidy scores of the ZIP zinc transporter genes across many epithelial types of cancer (compared to the established cancer genes or the SLC transporter control group) are compatible with the AMTC hypothesis. While such a genomic analysis alone cannot provide conclusive evidence for the proposed role of different ZIP genes as cancer driving genes, it nevertheless points in this direction.

**Figure 4:**
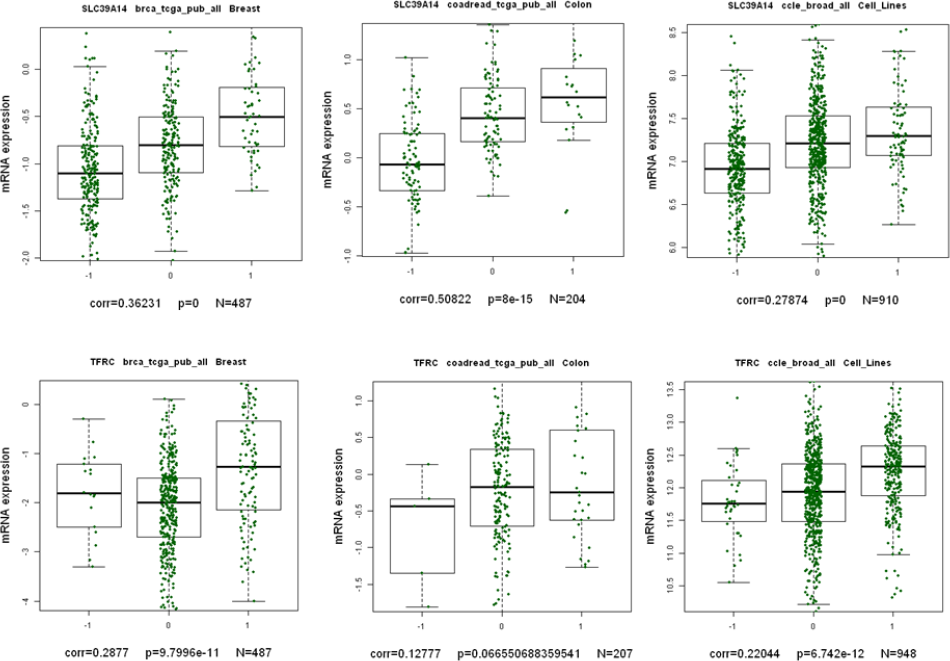
Gene dosage effects observed for two different metal transporter genes in different tumor types: Zinc transporter *SLC39A14* (upper row) and the iron transporter *TFRC* (lower row). A strong candidate driver gene can be expected to be hemizygously lost at high frequencies (one copy, termed “-1”, e.g. *SLC39A14*) or gained (three copies, termed “1”, e.g. *TFRC*) and hereby change the level of mRNA expresssion accordingly. For the analysis shown in Table 1, only results from genes that show a dosage effect in a specific type of cancer were taken into account. In this way, genes that only show high frequency changes but no gene dosage effect in a certain tumor type (possibly because they are poorly expressed in a certain tissue) were down-weighted. Gene dosage was defined as significant Pearson correlation [p<0.5, corr>0.1] between mRNA expression and copy number as indicated. Hence, of the six correlations above, only *TFRC* in colon cancer was not significantly correlated. Data was downloaded through the cBioPortal for Cancer Genomics ^105,106^ using the CGDS-R library.

The partial functional redundancy of dozens of metal transporters may possibly explain part of the observed strong differences in aneuploidy patterns between cancer types and subtypes resulting in similar - yet not identical - phenotypes. For example, different combinations of hemizygously gained and lost chromosome arms containing different metal transporters may help explain the characteristic aneuploidy patterns observed in different breast cancer subtypes. The general feasibility of such a classifying approach based on aneuploidy patterns was demonstrated ^109^ and visualized in Curtis et al. ^110^. A limited set of driver genes such as the group of metal transporters may thus represent a noteworthy alternative view to the continuum of many different driver genes acting through haploinsufficiency and triplosensitivity that was recently proposed ^111^.

Over the past decades, the long-neglected concept of aneuploidy and large chromosomal gains and losses as early driving force in cancer development seems to be regaining momentum ^111–113^, which calls for thorough statistical analyses of these non-focal events and for the identification of potentially overlooked functional groups of genes that, by copy number alterations and gene dosage effects, could contribute to carcinogenesis. We note that our model is significantly different from contributions by Peter Duesberg ^26^ in the sense that it is not the up- or downregulation of thousands of neighboring genes due to aneuploidy that is being suggested to be causal for cancer. It is also different from a recent study by Davoli *et al*. ^111^ who found strong correlations between the aberration frequencies of chromosome arms and different statistical measures of large numbers of putative tumor suppressor and oncogenes on these chromosomes, which was interpreted as indication that the cumulative effect of these genes may drive the observed aneuploidy patterns. According to our model, aneuploidy is also a causal factor for carcinogenesis, but aneuploidy evolves through the gene dosage effects of only a few causal genes, in particular metal transporter genes, whereas the numerous hitchhiking neighboring genes may, rather as a side effect, add variability to individual cancer clones.

## Trace elements and the hallmarks of cancer

Over the last decades, several now well-established hallmarks of cancer have been identified ^1,2^. These cancer hallmarks comprise biological capabilities acquired during the multistep development of human tumors and include sustained proliferative signaling, evading anti-proliferative growth suppressors, resisting cell death, enabling replicative immortality, inducing angiogenesis, and activating invasion and metastasis ^1^. Further, deregulating energy metabolism and evading immune destruction have been described as two emerging hallmarks of cancer cells, while genome instability and inflammation are now accepted as underlying enabling characteristics for cancer development ^2^. In the following paragraphs we would like to discuss how deregulated trace element homeostasis can impact these established hallmarks and enabling characteristics, and would like to speculate whether their acquisition could even be a response to imbalances in zinc and copper metabolism early during cancer development.

Trace elements are essential for the normal function of cells. Zinc, an essential trace element micronutrient is important in a wide range of cellular functions. It is a cofactor for cellular enzymes, and a structural component of many cellular proteins ^114^. Analysis of the human proteome indicates that zinc is a component of approximately 10% of human proteins ^115,116^. Thus, cellular zinc and zinc deregulation have the potential to modify the function of many important proteins. In addition to its role as a structural and catalytic factor, zinc also functions as an intracellular signaling molecule for differentiation, apoptosis, proliferation and cell cycle regulation. Hence, due to its ubiquitous presence and function in cells, zinc plays direct and indirect roles in the manifestation of the hallmarks of cancer identified by Hanahan and Weinberg ^2^. It is important to emphasize that we do not suggest that alterations in the homeostatic regulation of trace elements are necessarily involved with each of the cancer hallmarks. However, in the following paragraphs we argue that cells that have undergone imbalances in trace element metabolism may under certain conditions have a survival advantage, which can be attributed at least in parts to the hallmarks. Most of this discussion is focused on zinc, mainly because zinc is one of the best-studied of the essential metals and appears to be involved in many more regulatory cellular activities than other metals. However, one should bear in mind that imbalances in the levels of any one trace element could also affect the balance of other metals and thereby have broader impact on metal homeostasis and cellular function.

As for the number of identified mutations that affect the hallmark capabilities of a tumor, a recent study on lung adenocarcinoma ^117^ made an effort to map the identified driver mutations (point mutations and focal amplifications and deletions) to the extended hallmarks ^2^. Maybe surprisingly, none of the tumor samples showed mutations in all of the hallmarks, 38% of samples showed mutations in three or fewer hallmarks and 15% of samples did not show a single hallmark mutation, which was interpreted as indication that important driver mutations were still not identified ^117^. Likewise, it seems possible that certain driver mutations concurrently affect multiple hallmark capabilities. This would be congruent with theoretical expectations of relatively few driver genes per tumor. For example, the analysis of resequencing data from different tumors ^118^ in a Wright-Fisher evolutionary model resulted in an expected low number of 2 – 20 driver genes ^119^, concordant with the 5–8 events inferred from mathematical studies on age distributions at cancer onset ^120,121^. Possibly, these low numbers of identified and expected driver genes can be reconciled with the ten extended hallmarks in the framework of the AMTC hypothesis: a limited number of arm-level mutations may affect intracellular trace element concentrations in a cell and may hereby, in combination with additional mutations in established cancer genes, affect essentially all hallmark characteristics.

### Sustained proliferative signaling

Zinc is an important player in intracellular signaling processes in diverse cell types ^122^. An estimated 300 enzymes require zinc ^123^ and zinc itself may act as a secondary messenger or signaling ion, both in normal and malignant cells^124^. Zinc seems to affect the proliferative Ras signaling pathway, as shown in the model organism *C. elegans* ^125^, possibly through EGFR and Src as observed in different cell cultures ^126,127^. Zinc also has an effect on the levels of diverse tyrosine kinase receptors including ErbB2 with its role in cell proliferation ^128^. Also of note, it was shown that RNA Polymerase I may act as a last zinc reservoir under zinc-deficient conditions and hereby may couple proliferation with zinc homeostasis ^129^.

As described above, the tumor zinc level is consistently elevated in some cancers (e.g., breast) and consistently decreased in others (e.g., prostate, pancreatic, liver) ^130–132^. Thus, effects of zinc on proliferation and apoptosis are likely cell type specific and may differ in normal versus malignant cells. Moreover, the heterogeneity of tumors and the actual cellular level of zinc in normal and malignant components of the tumor add to the difficulty of defining the specific role of zinc in various cancers. Zinc has been shown to alter the PI3K/AKT pathway in MCF-7 cells, B-cells and MC3T3 ^133–135^. Zinc is required for normal cell cycle progression and is essential for transition from G1 to S phase and for DNA synthesis ^136–138^. These effects of zinc result in activation of proliferation and survival pathways. MAPK, ERK1/2, JNK and various tyrosine kinase signaling pathways are also activated by zinc ^126,139,140^. These reported effects of zinc are involved in proliferation signaling in normal and cancer cells. However, due to the large complexity of downstream effects of deregulated zinc homeostasis, exactly how zinc levels correlate with proliferative signaling in different cancer cells remains to be firmly established.

### Resisting cell death

Zinc is reported to both induce apoptosis and to protect against apoptosis in cells ^141,142^. Thus, the apoptotic effects of zinc are equally complex, inconsistent and controversial. The effects of zinc on apoptosis are studied mainly by exposing cells to extra amounts of zinc (in many studies unphysiological levels), by decreasing zinc in the medium to create zinc deficiency, and by treating cells with permeable chelators to deplete intracellular zinc. To address fully the issue of zinc and apoptosis, one must appreciate the complex biochemistry and physiology of zinc in biological systems and the various intracellular pools of zinc ^143–145^ and the experimental conditions of the reported studies. In interstitial fluid, the concentration of zinc is 2–5 µM, while the free zinc ion concentration is in the nM range ^146^. Intracellular zinc concentration is in the range of 100 to 500 µM. However, intracellular zinc exists in at least three intracellular pools. The vast majority (∼90%) is bound to cellular proteins and is not an available reactive pool. Zinc bound to low molecular weight ligands (amino acids, citrate, metallothionein) with low binding affinities and free zinc ion represent the reactive pool ^147^. The concentration of free zinc ion is in the fM-pM range and is not a physiologically relevant reactive pool of zinc ^148^. Thus, the complex physiology and biochemistry of zinc homeostasis requires special attention in the design and control of zinc levels, and in the interpretation of experimental zinc studies.

Nevertheless, zinc seems to play important roles in the determination and regulation of cell death by apoptosis ^58,149–153^. Zinc can have direct effects on the nucleus, mitochondria and protein factors to induce or inhibit apoptosis ^154–157^. Zinc can also have effects on pathways that regulate apoptosis ^158–160^. For example, the modulation of intra- and extracellular zinc was found sufficient to induce metabolic changes and apoptosis in different breast cancer cell lines^161^. While the mechanisms, cell specificity and pathways associated with this role of zinc are diverse and complex, the important consideration here is that deregulation of zinc homeostasis is potentially an important factor in the progression of most, if not all cancers.

### Evading growth suppressors

Transformed, neoplastic cells acquire the capability to evade growth suppressor effects that regulate the proliferation of normal cells. In addition to its role in modulating factors and pathways involved in proliferation and apoptosis, zinc is also essential to several proteins involved in DNA replication, DNA repair and the cellular response to DNA damage ^162–165^. Zinc is integral to many transcription factors that regulate key cellular functions. p53 is a tumor suppressor transcription factor that is implicated in development and progression of many cancers. It is a prototypical tumor suppressor, which is a component of many coordinated pathways leading to cell death in response to intracellular signals of cellular stress or injury ^166–169^. p53 is a zinc binding transcription factor whose activity is lost through loss of expression, decreased activation or genetic mutations ^85,170^. Zinc is reported to play a role in the mechanisms of p53 loss involving expression and activation particularly ^171–173^. Thus, deregulation of zinc homeostasis leading to changes in cellular zinc level will alter the ability of cells to respond to cellular stresses and DNA damage and may add a selection advantage for transformed cells to survive. Therefore, the deregulation of zinc homeostasis is a potentially important phenomenon in the acquisition of this hallmark of cancer.

Few studies on a direct involvement of zinc with retinoblastoma (RB) tumor suppressor pathways have been reported. However, the documented role of zinc in the activities of various phosphatases opens the potential for zinc to have a significant effect on the activity of RB tumor suppressor activity. In HL-60 and U-937 leukemic cell lines, apoptosis-specific RB dephosphorylation was inhibited by calyculin, A (a serine/threonine phosphatase inhibitor) but not by zinc ^174^. In contrast, hyperphosphorylated forms of RB were detected in zinc deficient esophagi^175^. Zinc is an inhibitor of phosphatases ^176,177^ and therefore, alterations in cellular zinc homeostasis could have effects on RB phosphorylation and its interaction with various RB binding proteins ^178^. However, at this point, there is no consensus and a role for metal dyshomeostasis in RB tumor suppressor function is speculation and awaits further exploration.

### Inducing angiogenesis

A supply of oxygen and nutrients and the removal of metabolic waste are essential for the survival of cells, and organs. The efficient supply of oxygen and nutrients and the removal of waste depend on blood vessels. Tumors can grow to limited size without blood vessels, however, for tumors to continue to grow and develop they require an established blood supply. New vessels develop from pre-existing vessels through branching and from the differentiation of endothelia cell precursors through the process of angiogenesis ^179^. Thus, angiogenesis is among the early events in the progression of all solid tumors. New vessel growth requires a change in the balance of angiogenesis activators and inhibitors in favor of angiogenesis ^180,181^. Essential metals modulate the activities of many of the angiogenic activators and inhibitors, thus changes in metal homeostasis may play a role in inducing angiogenic pathways.

Zinc and copper contribute to the regulation of the balance of angiogenesis activators and inhibitors. These metals function in the regulation of the signaling pathways leading to angiogenesis and the activity of key factors that are involved in the process. The angiogenic effect of copper has been known for decades ^182,183^. More recently, massive intra- and extracellular copper movements during angiogenesis were observed ^184^. Hypoxia-inducible factor 1 (HIF-1) is a transcription factor that is specifically activated during hypoxia and that regulates cancer related genes including those involved in angiogenesis e.g., VEGF and basic fibroblast growth factor (bFGF). Cobalt mimics the activation of HIF-1 and induces changes in gene expression through HIF-1 transcriptional activation similar to those caused by hypoxia ^185^. Zinc inhibits cobalt-induced changes in gene expression and reactivates certain drug-induced expression of proapoptotic genes ^185^. Thus, the hypoxia pathway and its regulation of angiogenesis are influenced by cellular zinc. NF-kappa B activates antiapoptotic genes, inflammatory cytokines, EGFR and VEGF, leading to increased proliferation and increased angiogenesis in many cancers. Zinc reportedly inhibits NF-kappa B through induction of A-20 and has beneficial effects on tumors by decreasing angiogenesis and induction of apoptosis ^186^. Copper also plays an important role in the early stages of angiogenesis ^187^. Copper is necessary for endothelial cell activation, it stimulates proliferation and migration. Copper activates VEGF, bFGF, TNF-α and interleukin-1 ^188^. Copper deficiency created either by diet or chelation blocks angiogenesis and downregulates the activity of VEGF, TNFα and IL-1.

Overexpression of ZIP4 caused a significant increase in expression of NRP-1, VEGF, MMP-2 and MMP-9 in both pancreatic cancer cell lines and xenografts presumably via altered intracellular zinc; however, zinc levels were not reported in this study ^56^. On the other hand, silencing of ZIP4 by RNAi decreased expression of NRP-1 and VEGF in pancreatic xenografts ^189^. In contrast, other studies report that the zinc level and the expression of ZIP3 are decreased in pancreatic cancer ^132^. While the effects of zinc on pancreatic cancer growth are controversial, several studies support a role for the deregulation of zinc and copper homeostasis in the angiogenic process.

### Enabling replicative immortality

The unlimited proliferative potential is a key characteristic of cancer cells. During the course of cancer development, cells acquire the capability to activate mechanisms that allow them to evade replicative mortality. Few studies have focused on a potential role of zinc and zinc level imbalances in this process. However, some links between zinc and replicative immortality exist, suggesting that deregulated zinc homeostasis and cancer cell immortality are not completely unrelated phenomena. For instance, it was recently reported that prostate cancer cells in their normal state are insensitive to zinc-induced senescence, a defect that could only be bypassed by additional infliction of DNA damage through ionizing radiation ^190^. Possibly, low zinc levels may also contribute to the de-differentiation of cells, e.g. it was observed that zinc is crucial in the differentiation of certain cell types ^191,192^. Beside zinc itself, this could include the transcription factor KLF4, which is known both as iPS stem cell factor in the induction of pluripotent stem cells ^193^ and as a regulator of the zinc transporter ZIP4 (SLC39A4) ^194,195^. On the other hand, telomerase activation by zinc and other metals was reported in human renal cell carcinoma and prostatic cancer cell lines ^196^, indicating that high zinc levels can also positively affect replicative immortality of cancer cells. Finally, zinc is a cofactor of several proteins present at telomeres. It is thus conceivable that changes in zinc levels might also entail deregulated telomere maintenance and telomerase function and thereby impact replicative immortality.

### Activating invasion and metastasis

Invasion and metastasis is a complex process that involves detachment of cells from the extracellular matrix, intravasations, resistance to apoptosis and survival in the lymphatic and systemic circulation, extravasation, reattachment in the new extracellular matrix environment and formation of small foci of cancer cells (colonization) leading ultimately to micrometastatic lesion. The major regulators of this complex process are mostly unknown. However, recent research suggests that transition to a more mesenchymal phenotype termed the epithelial-mesenchymal transition (EMT) is required for invasion and metastasis. Recently identified transcription factors regulate the EMT during embryogenesis and likely play a similar role in cancer invasion and metastasis. Expression of some of these transcription factors is regulated by intracellular zinc signals. ZIP6 has been linked to EMT through a STAT3 mechanism ^133,197^. During embryonic development in Zebrafish, STAT3 activates expression of ZIP6, which increases the nuclear translocation of Snail, which then represses the expression of E-cadherin and increases cell migration. Similar to Zebrafish development, Snail and STAT3 have been reported to have roles in cancer metastasis ^62,198^. Thus, alterations in the expression of zinc transporters and the presumed resulting changes in cellular zinc may provide an advantage for cells that have undergone additional genomic neoplastic changes.

Colonization can be separated from invasion as exemplified by squamous cell carcinomas that invade adjacent tissue, but rarely metastasize. While the EMT appears to be essential for the invasive phase, metalloproteinases (MMPs), a family of zinc dependent endopeptidases that degrade various components of the extracellular matrix, play an importation role in colonization. The actions of MMPs not only degrade the structure of the matrix but release growth factors and chemoattractants that recruit inflammatory cells to the invasions site ^199–201^. The growth factors and the activities of the inflammatory cells increase attachment of the cancer cells and resistance to apoptosis that allow the invading cancer cells to colonize. While there is little direct evidence for altered cellular zinc levels in the regulation of MMP expression and activation, the altered expression of various members of the zinc transporter families has been implicated in altered MMP expression in some cancers ^189,202^.

### Deregulated energy metabolism

Re-programming of energy metabolism is a widespread characteristic of cancer cells. The altered energy metabolism of cancer cells was initially recognized several decades ago by Otto Warburg ^203–205^ who noted that even in the presence of oxygen, cancer cell energy production is largely through glycolysis (aerobic glycolysis). Increased aerobic glycolysis results from upregulation of glucose transporters and increased utilization of glucose by the glycolytic pathway. This widespread characteristic of cancer cell metabolic re-programming is the basis for noninvasive visualization of tumors through positron emission tomography (PET), which depends on the uptake of a radiolabeled analog of glucose. Some links between zinc and glucose metabolism have been described, e.g. the role of ZIP7 in glycemic control in skeletal muscle ^206^ and the role of zinc and a polymorphism in the zinc transporter ZnT8 (SLC30A8) in diabetes ^207,208^. Although a general role for alterations in the cellular level of trace metals in energy metabolism re-programming is not well established for most cancers, in prostate cancer altered zinc accumulation is an important metabolic transformation.

Normal prostate epithelial cells are zinc-accumulating cells and contain high levels of cellular zinc ^144,209–213^. The high cellular level of zinc is required for the prostate function of citrate accumulation, which is a major component of prostatic fluid. Citrate accumulation is the result of limited citrate oxidation by prostate epithelial cells caused by zinc inhibition of mitochondrial aconitase. Malignant prostate cells, on the other hand, have lost the ability to accumulate zinc and thus contain significantly lower levels of citrate and zinc. Thus, the metabolic pathway of normal citrate producing prostate cells is similar to the metabolism of tumor cells; and the metabolism of the malignant prostate cells is similar to that of the normal mammalian cells. While there is no established relationship between energy metabolism re-programming and cellular trace metals for most cancers, the potential effect of an altered trace metal metabolism on energy metabolism is exemplified by the metabolic transformation in prostate malignancy and presents the potential for an association of altered cellular trace metals deregulation and energy metabolism.

### Evading immune destruction

The development of solid tumors in the presence of a competent immune system suggests that cancer cells manage to either evade immune surveillance or somehow disable the functions of immune cells. Thus, evading immune destruction is an emerging hallmark of cancer. At present there is little strong evidence supporting a direct role for altered zinc (or other metal) levels in tumor cell evasion of immune destruction; however, there is strong evidence for the role of Zn in immune system responses. Thus, it is tempting to speculate that cancer cells are selected for or evolve alterations in zinc homeostatic mechanisms that evade or disable the immune system and appropriate immune responses. Many early studies ^214–216^ have established that zinc is essential for recognition of antigens processed and presented by the proteins of the MHC class I and II to T cells and NK cells. In addition, zinc reportedly inhibits the trafficking of MHC vesicles to the plasma membrane from the perinuclear region. Thus, the fact that many cancers develop in the presence of decreased intracellular zinc raises the interesting possibility that the decreased cellular zinc may contribute to the evasion of the immune system by cancer cells. There is some evidence that tumors may have the ability to disable immune system signaling. A20, a cytoplasmic protein, which modulates immune responses, inhibits TNF-mediated cell death and NFkB activation and is regulated by zinc ^217–219^. A20 binds to TNF-receptor associated factor and inhibits the IKK-alpha NFkB pathway, thus protecting tumor cells against the actions of cytokines of the immune system. In view of a limited number of studies focusing on zinc and evading immune destruction, additional investigation of the alterations in metal homeostasis associated with many cancers and evasion of the immune surveillance by cancer cells is warranted.

### Genome instability

Zinc is an essential cofactor and structural component of various proteins involved in the surveillance of genome integrity. These proteins often contain zinc finger motifs that function as DNA binding modules; prominent examples include the tumor suppressor protein p53 (see above) and the DNA repair proteins OGG1, APE, and AP1, as well as poly(ADP-ribose) polymerase 1 (PARP1) ^190^. While zinc binding is a prerequisite for normal functioning of these enzymes, to what extent deviations in intracellular zinc concentrations affect their roles in DNA damage signaling and repair has remained ambiguous. In contrast, several studies have shown convincingly that zinc deficiency can lead to oxidative stress ^220–226^. It is thus possible that the accumulation of DNA damage as well as the deregulation of DNA repair protein expression observed under conditions of zinc deficiency largely reflect changes in the cellular redox status. Given the tumor barrier function of DNA repair mechanisms and genome stability, elevated levels of oxidative stress in response to reduced intracellular zinc concentrations could provide one mechanism how cells reach the threshold to become genomically unstable and cancer prone.

### Inflammation

Zinc is deregulated in diverse inflammatory processes ^227^ including inflammation of the skin ^228^, airway epithelium ^229^, bowel tract ^230^ and in response to endotoxin exposure ^231^ and allergic reactions ^232^. In response to viral or bacterial pathogens, trace element concentrations are also deregulated (Appendix). On the protein level, interleukin-6 contributes to hypozincemia during the acute phase response by its effect on the zinc transporter ZIP14 (SLC39A14) ^233^. The zinc transporter ZIP8 (SLC39A8) regulates host defense against bacterial infection through its interaction with the pro-inflammatory transcription factor NFkB ^234^. Further, zinc deficiency seems to induce interleukin-1ß ^235^ which in return contributes to the control of metallothionein expression ^236^ and SLC39A14 ^237^. In summary, zinc homeostasis and the immune system are closely linked ^238^ and trace elements play important albeit not entirely understood roles during inflammatory processes.

Collectively, deregulated zinc homeostasis may directly or indirectly impact on several of the major hallmarks of cancer, most prominently on proliferative and anti-proliferative signaling, apoptosis, inflammation and genome instability. While the underlying mechanisms and the overall contribution for tumor development and progression are complex and incompletely understood, the established links provide an entry point for future research to elucidate how trace element homeostasis and aberrations thereof affect cancer-relevant biological pathways.

## Outlook and possible implications for cancer prevention, diagnosis and therapy

First of all, and regardless of the AMTC hypothesis, currently there exists a huge potential of lessening the burden of cancer by avoiding lifestyle risk factors as well as through early diagnosis and improved therapy.

While the known **prevention efforts** related to the avoidance of risk factors (smoking, obesity, etc.) remain, a new focus may be on a healthy trace element profile of the individual. One way to lower cancer risk might be to maintain a healthy trace element homeostasis. However, this balance is complex and currently poorly understood. Therefore, the best way to achieve a healthy trace element homeostasis may be through a healthier lifestyle rather than through dietary supplements.

For **diagnosis and monitoring**, it may be helpful to combine genomic tools such as CGH analysis with ionomic tools. A personal trace element profile for each patient that would include trace element measurements from tumor biopsies and serum may turn out helpful. For example, zinc/copper ratios and the estimation of trace element concentrations in hair samples may be of value ^239^. Such trace element profiles and mutational analyses might also have prognostic value and guide treatment decisions for or against certain cancer therapies. Moreover, as part of the emerging field of personalized medicine, circulating cells or tumor biopsies could be interrogated for genomic patterns of gain and loss that specifically affect genes related to copper and zinc homeostasis and could be used as a diagnostic marker for the presence of tumors and even of tumor subtypes. Optimistically, combinations of different gains and losses including metal transporters may help explain different characteristics of tumors and thus be used in diagnosis. Mutational analyses have recently been done in circulating cells in the blood stream by resequencing ^240^ and array CGH ^241^ an approach which may simplify the identification of patient-specific driver genes in the future. These advances may certainly profit from single cell sequencing ^242^. Last but not least, bioimaging techniques for cancer diagnosis and imaging ^243^ may improve future cancer diagnosis.

**Therapy** may on the long term have an additional focus to restore trace element concentrations. Pharmaceutical or dietary interventions and chemotherapeutic agents designed to restore the balance of intracellular trace element concentrations might become a focus of cancer therapy. One paradigm might be to change the tumor microenvironment in such a way that the tumor cells lose their putative selective advantage over normal cells in a tissue, e.g. by suppressing the inflated copper concentrations and increasing the depressed zinc concentrations in a tumor. In this context it is interesting to note that several copper chelating agents are candidate drugs for antiangiogenic therapy in cancer ^244–246^, and in particular, tetrathiomolybdate has been successful in several clinical trials ^247,248^. Likewise interesting, some authors have discussed the cytotoxic and tumor-suppressive properties of zinc ^249,250^, and intra-tumoral zinc-acetate injection was shown to halt tumor growth in a mouse xenograft model of prostate cancer ^251^. Other potential anti-cancer agents which alter zinc homeostasis have also been reported ^252^. Moreover, by combining the copper chelator trientine with platinum-based drugs in a pilot study on high-grade epithelial ovarian cancer it seemed possible to partially overcome drug resistance ^253^. If possible links between trace element homeostasis and the functional mechanisms of resveratrol or cisplatin (Appendix) turn out to be vital, there may be ample opportunities to improve our understanding of commonly used cancer drugs and, in the long run, hopefully improve therapy. Further, dosing schemes of different anti-cancer drugs may improve from considering life history theory ^254^, e.g. with the goal of increasing the time before relapse. In summary, modulation of trace element homeostasis through ion chelators and through targeting key regulators or metal transporters might be a promising way for future research in cancer therapy.

In **basic research**, to understand the role of each transporter in different organelles and in a supposed protein-protein interaction with metalloenzymes and other metal binding molecules would be of utmost interest, both in relation and beyond its function of changing intracellular element concentrations. Zinc fluorescent probes ^255^ as well as zinc chelators ^256^ may play important roles in this regard. As a very general outline, quantitative models of trace element homeostasis over different spatial (molecules to organelles to cells to tissues to species to ecosystems) and temporal (from protein kinetic to circadian to seasonal) timescales are needed. Zinc as an example may play a role in diseases well beyond cancer, including different neurological diseases, depression, diabetes, diarrhea, sepsis, atherosclerosis, stroke and eye diseases among others ^257,258^. Given the wide spectrum of biological functions of essential trace elements, the knowledge gained from such efforts may eventually feed back into other areas of biology.

## Conclusions

Based on a wealth of studies from the fields of trace element and cancer biology, we have used evolutionary genomics to build a hypothesis related to carcinogenesis. We emphasize that we do not present here direct experimental evidence for our main hypothesis that chromosomal gains and losses of metal transporters have a causal role in carcinogenesis. Instead we point in different directions how this could be tested (Text Box B). If confirmed, the disruption of zinc, copper and iron homeostasis would emerge as a unifying theme across diverse types of cancer with implications for its prevention, diagnosis and treatment. We hope that this article will encourage studies in different areas of biology to rigorously test these ideas in the future.

TEXT BOX A: Glossary

Definitions and concepts may vary, overlap in meaning, and can be subject to change as our understanding of cancer development and progression advances.

**Amplification:** Genetic alteration resulting in the duplication or amplification of a genomic region; can cause *gene dosage effects*. Amplifications of whole chromosomes result in *aneuploidy*.

**Aneuploidy:** Numerical *chromosomal aberration* characterized by any deviation from normal chromosome number, i.e. having extra or missing chromosomes or chromosome arms; antonym to *euploidy*. Aneuploidy can cause *gene dosage effects*.

**Chromosomal aberration:** Any irregularity or abnormality of chromosome structure or number. Structural chromosomal aberrations include *amplifications*, *translocations*, inversions, deletions and insertions; numerical chromosomal aberrations result in *aneuploidy*.

**Chromosomal gain:** Genetic alteration resulting in the duplication of a large chromosomal region or of an entire chromosome.

**Chromosomal loss:** Genetic alteration resulting in the deletion of a large chromosomal region or of an entire chromosome. Leads to *loss of heterozygosity*.

**Chromothripsis:** Genetic alteration that is characterized by the acquisition of massive and complex genomic rearrangements in a single catastrophic event involving the almost simultaneous fragmentation and reassembly of chromosomes.

**Clonal evolution:** A process of Darwinian natural selection, by which cancer cells change their genotype and phenotype over time through a reiterative process of genetic diversification, clonal expansion and clonal selection.

**Driver gene**: Gene that, when mutated, provides a selective growth advantage and promotes cancer development.

**Driver mutation:** Cancer-specific genetic alteration of a *driver gene* that results in a selective growth advantage and facilitates cancer development.

**Euploidy:** State of normal chromosome number, i.e. an integral multiple of the haploid number of chromosomes.

**Focal amplification:** *Amplifications* that, as opposed to amplifications of whole chromosomes or chromosome arms, affect relatively small regions of genomic DNA.

**Focal deletion:** Homozygous deletion that, as opposed to deletion of whole chromosomes or chromosome arms, affect relatively small regions of genomic DNA.

**Gene dosage effect:** Condition in which altered gene copy number correlates with gene function, i.e. an increase in gene dosage causes higher levels of gene product and increased function, whereas a decrease in gene dosage causes lower levels of gene product and reduced function.

**Haploinsufficiency**: A specific *gene dosage effect* in which a single copy of a particular gene is insufficient to support the function normally provided by the two alleles in a diploid genome.

**Homeostasis of trace elements:** A process that maintains the concentrations of trace elements stable and in balance to interior and exterior influences.

**Homozygous deletion:** Deletion of both copies at a specific genomic locus in a diploid genome.

**Loss of heterozygosity (LOH):** Loss of one copy at a specific genomic locus in a diploid genome (hemizygous loss). LOH can cause *gene dosage effects* and result in complete loss of gene function if the second, remaining allele is mutated. *Tumor suppressor genes* are often affected by loss of heterozygosity.

**Oncogene:** Gene that, when abnormally activated by mutation confers a selective growth advantage and promotes cancer development.

**Passenger gene:** Gene that is frequently mutated in cancer cells without providing a selective growth advantage.

**Passenger mutation:** Cancer-specific genetic alteration of a *passenger gene* or cancer gene that does not result in a selective growth advantage.

**Point mutation:** Mutational event of a single DNA base substitution.

**Positive selection:** Process by which one phenotype is favored over another phenotype, i.e. profits from a growth advantage, and thus is selected for.

**Somatic copy-number alterations (SCNA):** SCNAs can be divided into arm-level SCNAs (*chromosomal gains and losses*) and focal SCNAs (*focal amplifications and deletions*).

**Translocation:** *Chromosomal aberration* that is characterized by a rearrangement of segments between non-homologous chromosomes.

**Triplosensitivity**: A specific *gene dosage effect* in which three copies of a particular gene lead to aberrant gene function.

**Tumor suppressor gene:** Gene that, when abnormally inactivated by mutation, confers a selective growth advantage and promotes cancer development.

TEXT BOX B Testing the AMTC hypothesis

The AMTC hypothesis is based mainly on a tentative analysis of genomics data, which suggests that gains and losses of metal transporter genes occur frequently, correlate well with transporter gene expression levels, and may underlie the observed aneuploidy in cancer. From this starting point, a number of informative experimental and statistical approaches can be outlined:

1. Experimental: To recapitulate hemizygous gains and losses of one or several candidate metal transporters in cell culture models that show 50% (loss) or 150% (gain) of normal gene expression and to characterize these models on the genomic, transcriptomic and ionomic level and with regard to cancer cell characteristics such as cell proliferation and invasiveness in order to test whether they are transformed towards “malignancy”.
2. Experimental: Similar experiments could be done with targeted gains and losses of entire chromosome arms, e.g. the introduction of additional copies of 1q and 8q in liver cell lines.
3. Experimental: Attempt to modify the trace element homeostasis in xenograft mouse models and hereby influence the course of carcinogenesis.
4. Experimental: Compare chromosomal gain and loss patterns, gene expression and trace element concentrations (if possible labile versus bound elements and in different cell compartments) in paired tissue samples (healthy versus tumor) and between individuals. If done with enough samples, one could test whether certain chromosomal gains and losses or combinations thereof are associated with differences in intracellular trace element concentrations, and in extension, whether they are associated with a response to anti-cancer or cancer-promoting compounds (cisplatin, estradiol, resveratrol, aspirin, etc.).
5. Bioinformatic analysis: Compare gain loss patterns of different cancer subtypes with their preferred tissue of metastases. Roughly following the seed and field hypothesis, one might expect that metastases occur preferentially in tissues that support the expression of the aberrant driver genes.
6. Experimental: Analyze normal healthy tissue and early cancerous lesions by single cell sequencing in order to learn about the background mutation burden in healthy tissue and early steps in carcinogenesis.
7. Experimental: Carefully compare chromosomal gains and losses with gene expression similar to the presented analysis but in a genome-wide manner and in many additional cancer types and subtypes.
8. Bioinformatic analysis: Compare chromosomal gain and loss patterns of candidate genes in diverse types of cancer with their mRNA expression pattern of candidate driver genes in matched normal tissue. As an additional criterion to identify driver genes in chromosome arm events, strong candidates should show a better correlation than passenger genes. In addition, driver genes should show a gene dosage effect in cancer samples. Prioritize genes by finding best correlation between mRNA expression in normal tissue against aberration frequency in the corresponding cancer.
9. Comparative oncogenomics: Compare CGH profiles between human, mouse, rat, dog and zebrafish cancers. Such approaches have been used test whether candidate loci are affected similarly in humans and model organisms. Specifically, it should be tested whether the proposed metal transporters are located in overlapping regions of recurrent chromosomal losses and gains (also see Supplemental Note D).
10. Experimental evolution: Raise cisplatin-resistant cells from sensitive mother clones by exposure to cisplatin and then test for chromosomal gains and losses. It would be expected that chromosomes with certain copper transporters are gained or lost preferentially. In addition, test whether e.g. the gain of 13q enhances cisplatin resistance through ATP7B overexpression. Such questions can be pursued in a framework of *in vitro* cancer evolution ^259^.
11. Experimental evolution: Expose non-cancerous, immortalized cell lines to zinc (copper, iron etc.) starvation and toxicity and later test for chromosomal gains and losses. It would be expected that certain chromosomes are gained or lost preferentially as a function of gene content, selection coefficient and generation time. For example, it might be expected that non-cancerous prostate, breast and liver cell lines would adapt to zinc deficient conditions by clonal expansion of mutated cells with certain chromosomal gains and losses. When replicated a sufficient number of experiments, the resulting genomes may recapitulate the aneuploidy patterns of different cancer types (e.g. prostate, breast and liver cancer).
12. Experimental: Test whether certain viral and bacterial infections change the trace element dynamics of infected cells, tissues and organisms.
13. Epidemiological: Analyze different measures of trace element status in population studies of cancer incidence and survival.

## Acknowledgments, online resources and author contributions

### Acknowledgments

We thank all Cancer Patients & Sample Donors that have indirectly contributed to this work. We apologize to the many authors that we failed to cite. We acknowledge the following scientists for helpful comments and corrections (in alphabetical order): Diego Hartasanchez, Fyodor Kondrashov, Leslie Costello and Lucas Carey. We thank Brandon Invergo for English correction and Verena Vogler for help with graphics. J.E. acknowledges the Volkswagenstiftung, the Arbeitsamt Waiblingen and the Servicio Público de Empleo Estatal (SEPE), M.A. the Novo Nordisk Foundation and the Danish Council for Independent Research for financial support. The authors declare that they have no conflict of interest.

We further acknowledge the following online resources:

http://cancer.sanger.ac.uk/cancergenome/projects/census/

http://www.intogen.org/

http://Progenetix.org

http://cran.r-project.org/web/packages/cgdsr

http://www.cbioportal.org

http://www.icgc.org/

http://cancergenome.nih.gov

https://confluence.broadinstitute.org/display/GDAC/Home

http://compbio.med.harvard.edu/metacghBrowser

http://www.ncbi.nlm.nih.gov/pubmed

http://scholar.google.com

http://www.wikipedia.org

### Author contributions

Developed the original AMTC hypothesis, analyzed the genomics data, and drafted the sections „Model of carcinogenesis” and “Possible implications of AMTC hypothesis” (JE). Drafted the section “Trace elements and the hallmarks of cancer” (RF, MA). Drafted the section “Testing the AMTC hypothesis”, and integrated and further developed the different sections (JE, MA). Provided the glossary (MA). All authors elaborated on the idea and discussed the manuscript.

## APPENDIX Possible implications of the AMTC hypothesis

The following is an attempt to describe possible implications of the AMTC hypothesis on different fields of research. While some points may be overly speculative, we hope to stimulate fruitful discussion and a possible re-evaluation and re-interpretation of existing studies.

### Chemotherapeutic agents and resistance

The platinum-based anti-cancer drug cisplatin is a DNA-damaging agent that causes DNA cross-links ^260^ and problems during DNA replication, which eventually result in cell death. Of note, cisplatin heavily relies on the cellular copper transport system by using different copper transporters for cellular entry, e.g. the copper transporter CTR1 (SLC31A1) ^261^. Biochemically, a methionine motif in CTR1 and possibly other copper transporters interacts with platinum-based drugs ^262,263^. In the copper transporter ATP7B, a Cys-X-X-Cys motif was found to be essential for cisplatin transport ^264^. Specifically, cisplatin uses the Cys-X-X-Cys sequence motives in ATP7B and mimics Cu in its transport characteristics ^265^. Further, it is known that cisplatin resistance can be mediated by the copper transporter ATP7B ^266,267^ as well as by the copper transporter ATP7A ^268,269^ and by the copper transporter CTR1 ^270^. Attenuation of ATP7B in different cell lines made different cisplatin-resistant cancer cell lines highly sensitive to cisplatin ^271,272^.

The emergence of drug resistant cell clones poses a difficult challenge in cancer therapy. Resistance is likely encoded in the resistant cellś genomes, and for some drugs, causal mutations could be identified, e.g. a point mutation in EGFR conferring resistance to gefitinib ^273^ or the amplification and overexpression by gene dosage of ERBB2 leading to tamoxifen resistance ^274^. For other drugs such as cisplatin, the resistance mechanism is less clear. Interestingly, cisplatin resistance is strongly associated with characteristic chromosomal gains and losses when compared to cisplatin-sensitive tumors ^275,276^. For example, the chromosome regions 1q21–22 and 13q12–14 seem recurrently gained in cisplatin resistant ovarian tumors ^277^ and intriguingly, we note that that these genomic region encode the zinc transporter SLC39A1 and the copper transporter ATP7B. Importantly, in evolutionary experiments it has been shown recently that yeast can quickly evolve multi-drug resistance both by point mutations and by chromosomal aberrations ^278^. A similar result was found in *Candida albicans* which acquired resistance against the anti-fungal drug fluconazole through the early appearance of aneuploidy ^279^. These observations lead to the question, whether chromosomal aberrations could also be a causal or contributing factor in the emergence of drug resistance in human cancers.

Taken together, the observed interactions of cisplatin with cellular copper transport and the recurrent chromosomal aberrations in cisplatin resistance raise the question, whether chromosomal gains or losses may confer resistance to platinum-based drugs by gene dosage effects of copper transporters (or antagonizing zinc transporters). Possibly, the characteristic enhanced uptake of copper in tumor cells (as shown by their increased copper levels) may lead to an enhanced uptake of cisplatin and hereby selectively lead cancer cells to apoptosis. Conversely, a cisplatin-resistant cell may shuffle cisplatin out of the cell, e.g. by overexpressing ATP7B. In other words, it seems possible that the potential of platinum-based compounds to specifically target cancer cells (as compared to non-cancer cells) could be at least partly related to preferred drug uptake by transformed cells with deregulated metal transporters.

Additional chemotherapeutic and preventive agents like resveratrol, curcumin, aspirin, artemisinin, or clioquinol should be systematically studied to test whether they interfere with trace element homeostasis or metal transporters. In this context it is interesting to note that the anti-cancer drug resveratrol at certain concentrations was already shown to increase the intracellular zinc levels in human prostate epithelial cells ^280^. Curcumin and resveratrol in combination may have a synergistic effect on zinc levels during lung carcinogenesis ^281^. Some weaker hints that there could be an interaction between zinc and aspririn also exist ^282,283^. As a side-note, the copper metabolism protein COMMD1 interacts with the anti-tumor peptide CIGB-552^284^ and its suppression correlates with an invasive tumor type ^285^. The metal cation-containing chemotherapeutic drug motexafin gadolinium has been shown to change zinc homeostasis and induce metallothionein and expression of the zinc transporer SLC30A1 ^286^. Of further note, certain metal ionophores have been shown to possess anti-cancer properties ^287,288^.

Copper-chelating agents such as penicillamine or alternatively, trientine or tetrathiomolybdate are used to treat Wilson’s disease, a hereditary disease characterized by copper overload of the liver. Interestingly, all of these agents, penicillamine ^245^, trientine ^289^ and tetrathiomolybdate ^188^ (e.g. ammonium tetrathiomolybdate or choline tetrathiomolybdate, ATN-224) show chemotherapeutic properties in cancer models or clinical trials. The investigational anti-cancer drug elesclomol is also a copper chelator. On this background, together with the observed increased concentrations of copper in cancer tissue and serum, the question has been raised whether copper chelation or copper deficiency may be a therapeutic strategy against cancer ^34,290–292^. While the mechanism of action of these copper-chelating agents is commonly attributed to their inhibitory effect on angiogenesis or reactive oxygen species, a complementary effect on cancer through the direct interference with trace element homeostasis should not be ruled out.

While all these reports strengthen the proposed link between zinc/copper homeostasis and chemotherapeutic drug treatment, it will be important to experimentally test possible molecular mechanisms underlying the observed effects.

### Metastasis

Intratumor heterogeneity including metastases in kidney cancer can be understood as a branching evolution pattern ^293^, leading to independently evolving metastatic clones that can be quite different from each other ^294^. The phenotypic and functional properties of metastasis have been described in the framework of cancer hallmarks, but their genetic basis is not entirely understood. Metastatic tumors are characterized by large numbers of point, focal and chromosomal mutations, which may be a side effect of impaired DNA repair and increased genomic instability. Alternatively, a subset of these mutations may be active drivers towards metastasis.

In the latter direction, statistically significant differences between aneuploidy patterns have been described in diverse combinations of different primary tumors and different sites of metastasis. In colorectal cancer to liver metastasis, gains at 8q23–24, 15q21–26, 19p and 20q and losses at 18q12–23 were observed more often in the metastatic than in the primary tumor ^295^. Notably, these regions encode the zinc transporters SLC39A4, SLC30A4, SLC39A3 and SLC39A6 (among many other proteins). However, this seems different from a similar study of colorectal cancer, that found common aberrations at 8p, 17p and 18q to predispose to metastases in liver ^296^. In gastric cancer to lymph node metastasis, gains of 20q12–13 and losses of 21q cen-21, 4q and 14q22-ter were identified ^297^. In renal cell carcinoma metastasis, gains of 1q21.3, 12q13.12, 12q13.3q14.1 and 20q11.21q13.2, and loss of 9p21.3p24.1 ^298^ containing zinc transporter genes *SLC39A1 SLC39A5* among many others. In pharyngeal carcinomas to lymph node metastasis, gains in 11q13 and 22q and losses of 18q were observed ^299^. In ocular tumors, the highest metastatic rate was found in a subgroup defined by the gain of 8q and losses of 3, 8p, and 16q ^300^. In breast cancer to brain metastasis, cytoband 11p15 was identified ^301^, with no candidate metal transporter (but *SLC39A13* nearby). Interestingly, lymph node metastasis in oral cancer was associated with loss of one copy around cytoband 13q14.3 ^302^, a region containing copper transporter *ATP7B*.

Excitingly, the introduction of human chromosome 8 leads to suppression of metastasis in rat tumor cells ^303^, a candidate region which was later mapped to the short arm of chromosome 8, more precisely the region 8p21-p12^304^. This would be consistent with the localization of human zinc transporter *SLC39A14* (8p21.3). Therefore, it may be asked whether this phenomenon could be driven by an increased expression of zinc transporter *SLC39A14*. Contrary to their experiment, *SLC39A14* is typically hemizygously lost in primary or metastatic tumor cells.

With regard to trace element concentrations, it was observed that zinc may have an influence on metastasis ^305^. Different types of metastases show deregulated trace element concentrations compared to surrounding normal tissue ^306,307^. As for metal transporters, ZIP10 is involved in metastasis of breast cancer and its knockdown led to decreased migratory behavior in breast cancer cell lines ^308^. Overexpression of SLC39A4 was linked to cancer aggressiveness in the pancreas ^56^ and liver ^58^. Alternatively, since *SLC39A4* and the oncogene *MYC* both locate to cytoband 8q24, *MYC* may be the relevant driver gene. Fittingly it was suggested that the prognostic signatures of primary tumors towards metastasis may reflect the activity of the *MYC* oncogene ^309^. Possibly, both genes could play a role, e.g. *SLC39A4* by chromosomal gain and *MYC* by focal amplification.

In the light of the “seed and field hypothesis” ^310,311^, the site where metastases spread is not arbitrary but related to a good match between certain properties of the circulating cancerous cell and its potential host tissue. May such metastatic behavior be related to zinc? Following the seed and field hypothesis and modifying it to the special case of trace elements, those “fields” (tissues or organs) that encourage the expression of a certain metal transporter would be a better environment for a particular “seed” (cancer clone) possessing the same transporter in a genomically amplified region compared to those fields in which this transporter does not have the right conditions. Such impact from the microenvironment among many other characteristics may include extracellular signals like interleukins or the trace element concentrations themselves. The tissue expression of a certain transporter is not directly relevant for the expression of that transporter in the cancerous cells, but to a limited extent it may possibly serve as a proxy for the right conditions. A possible example may be SLC39A14 which is induced by IL-6 in liver ^233^; the same region is often lost in colorectal liver metastases (see below). As a consequence, metastasis may occur more often in organs or tissues with a fitting expression pattern of the metal transporter in question.

In summary, we describe the observation of increased aneuploidy in metastatic tumors and a number of intriguing but so far not fully conclusive observations related to zinc. This leads us to the testable hypothesis that metastatic cell clones may be dominated by acquired chromosomal aberrations, which further disrupt the intracellular trace element concentrations and hereby, possibly following the seed and field hypothesis, facilitate the spread of metastases.

### Viral and bacterial infection, trace element homeostasis and cancer

It is well established that infection with certain bacteria and viruses increases the incidence of a number of cancer types. For example, cervical cancer is strongly associated with papillomavirus infections ^312^. Likewise, gastric cancer development is facilitated by the widespread bacterium *Helicobacter pylori* ^313^. In turn, the treatment of *H. pylori* seems to be efficient against gastric cancer ^314^. Burkitt’s lymphoma is associated with the Eppstein Barr Virus ^315^. Kaposi’s sarcoma in HIV infected individuals can be caused by a certain type of herpesvirus presumably through a mechanism termed “paracrine neoplasia” affecting cell proliferation and apoptosis ^316^. Additional links albeit less studied may exist between non-melanoma skin cancers and human papillomaviruses ^317^. Interestingly it is not entirely clear what biological mechanism may drive this phenomenon. Several mechanisms have been proposed to explain the role of viruses in the etiology of cancer: (i) viral oncogenes ^318^ (ii) functional inactivation of tumor suppressor genes ^319^ (iii) inflammatory processes ^320^ (iv) epigenetic mechanisms ^321^. As a bottom line, in all types of cancer associated with either bacterial or viral infections, inflammatory processes and the immune system may play a key role ^322,323^.

Here we ask the provocative question whether the disruption of trace element homeostasis during infection may be a complementary contributing force in these types of cancer? On the one hand, inflammatory processes during infection are partially mediated through zinc ^238^. On the other hand, following the concept of “nutritional immunity” ^324^, pathogens are often starved from iron, manganese or zinc as part of the immune defense. It is unclear however, whether this concept may apply to viruses as well. For example, may oncoviruses trigger local zinc deficiency in the host (nutritional immunity) and hereby facilitate the subsequent development of cancer? While there has been little research into the interplay between nutritional immunity and viral infection, the inflammatory response triggered by viruses likely profoundly impacts vertebrate metal levels (Eric Skaar, personal communication).

Bacterial and viral infections are intimately connected to perturbances in the host´s homeostasis of trace elements. For example, infections with *S. aureus* trigger a highly localized reduction of zinc and manganese in infected tissue^325^. *H. pylori* infections are associated with iron deficiency ^326^. Low zinc levels in gastric tissue were correlated with an increased degree of inflammation caused by *H. pylori* ^327^. In a genome-wide screen, H. pylori infection was shown to lead to differential expression of 38 genes in gastric tissue including genes involved in inflammation and several regulators of trace element homeostasis ^328^. Studies on oncoviruses have shown that in some cases, signaling via STAT3 ^329^ or the IL-6-STAT3 axis ^330^ was involved in tumorigenesis. These studies did not interrogate zinc concentrations, but other studies suggest a link between STAT3, IL-6 and zinc ^331,332^. Fittingly with an important role of trace elements in viral infections, it has been observed that metal ionophores, particularly zinc ionophores, exert a strong effect on infection and replication of different types of virus ^333–335^. Further, the human EVER proteins 1 and 2 (TMC6 and TMC8) have been found to interact with zinc and the zinc transporter SLC30A1 during papillomavirus infection ^336,337^. As a side note it has long been known that the relevant proteins E6 and E7 in papillomaviruses possess a Cys-X-X-Cys zinc-binding sequence motif ^338^ which is essential for host cell transformation ^339^.

In summary, in addition to the established link between pathogenic infections and cancer, there is a strong implication of trace elements in bacterial and viral infections. Considering the AMTC hypothesis, it therefore seems justified to speculate about an increased cancer risk through viral and bacterial infections, facilitated by their suspected temporal effects on tissue trace element concentrations which could provide an early selective force in carcinogenesis. However, additional and more direct experimental evidence is needed.

### Hormone dynamics

Certain subtypes of breast, ovarian and prostate cancer share a similar etiology related to sex hormones which was the rationale behind the decision to study them together in the Collaborative Oncological Gene-environment Study^340^. Indeed, several of the identified genetic risk loci are pleiotropic, acting in more than one cancer type ^340^. Interestingly, cell lines from other types of cancer such as colon cancer also respond to sex hormones and to the hormone-blocking tamoxifen ^341^. Different hormone receptors like the estrogen and progesterone receptors play a role in these interactions, but the functional mechanisms are not entirely understood.

A number of interactions between sex hormones and zinc or zinc transporters have been documented. For example, zinc and other metals activate estrogen receptor-a (ERa) in the human breast cancer cell line MCF-7 ^342^, estrogen decreases ZnT3 (SLC30A3) and synaptic zinc ^343^ and 17ß-estradiol influences metallothionein and zinc levels in squirrelfish ^344^. ZIP6 (SLC39A6) is regulated by estradiols ^345^, ZIP7 modifies the phenotype of a tamoxifen-resistant breast cancer cell line ^128^ and ZIP6 has a role in the epithelial-mesenchymal transition of breast cancer cells ^63^. Further, treatment of prostate cancer cells with prolactin and testosterone increased their zinc uptake, possibly through an increased activity of ZIP1 ^346^

Different subtypes of estrogen receptor positive (ER+) and negative (ER-) types of cancer have characteristic chromosomal aberration patterns ^347–351^,^352^, ^109^. Whether some of these patterns could be related to the presence of metal transporters according to the AMTC hypothesis, remains to be tested. Fitting with such a scenario is the observation that tamoxifen-resistant clones of a tamoxifen-sensitive MCF-7 breast cancer cell line showed a number of recurrent chromosomal aberrations that did not contain estrogen receptor coding regions ^353^ but which contained, among many other genes, a number of different zinc and copper transporter coding regions.

In summary, some important functional links exist between hormone and zinc transporter dynamics, but this leaves the question open whether aneuploidy patterns and their proposed effect on zinc transporter expression could exert an influence on hormone dependent characteristics of certain types of cancer.

### Cancer risk in obese populations

In epidemiological studies, a strong link between obesity and cancer risk has been observed ^354^ and different mechanisms related to insulin, adipokines and obesity-induced hypoxia have been suggested ^355^. However, the evidence seems inconclusive and the topic was highlighted in the recent “Provocative Questions Initiative” by the NIH ^356^. In the following we briefly discuss whether the effect of obesity on cancer risk may be reconciled with the AMTC hypothesis.

First and overall, there is a significant effect of obesity, age, gender and diet on serum zinc and other trace elements ^357,358^. Typically, serum zinc is reduced in obese people ^359,360^. While this relationship may be reversed in hair samples ^361^ and an opposite trend was observed in obese children ^362^, it is supported by results from obese transgenic (ob/ob) mice, which show reduced zinc and copper levels in diverse organs ^363^.

Second, it was observed that mice subjected to a carcinogen only developed cancer when fed a high fat diet but not a low fat diet, an effect which the authors attributed to the gut microbiota and especially to the effect of deoxycholic acid ^364^. In general, this result seems to be congruent with our notes on the evolutionary dynamics of cancer that propose a joint effect of mutations and selective forces as indispensable factors of carcinogenesis. In our interpretation, the putative selective force of local zinc deficiency would either be explained by the zinc-limiting effect of fat alone, or alternatively, by a more complex interaction that could involve inflammation or the possible relationship between deoxycholic acid and zinc ^365,366^. Of interest, there exists a possible regulatory role of zinc on leptin levels ^367^ and there are known links between leptin and obesity ^368,369^ as well as between leptin and cancer ^370^. So it would be interesting to test whether there exist causal links between leptin metabolism and cancer through zinc.

Third, we note that the occasionally observed elevated levels of oxidative damage surrounding tumors may be a direct result of low zinc concentrations rather than an effect of the tumors. Support for this idea comes from a study of rat liver which suggested that “increased hepatic microsomal lipid peroxidation was associated with zinc deficiency whether using in vivo or in vitro indices of measurement” ^371^. The results of a study in rat testes ^226^ point in a similar direction.

In conclusion, it seems possible to explain the link between obesity and cancer in a way that is congruent with the AMTC hypothesis. Obesity may lead to local zinc deficiency, which in turn (i) may increase oxidative damage and (ii) may contribute to a microenvironment acting as a selective force that promotes carcinogenesis. However, direct experimental tests of this interpretation are needed before we can be confident of its usefulness.

### Cancer risk in the elderly population

To some extent many cancer types are more common in elderly people ^372,373^ a finding which may be consistent with accumulation of DNA damage over time and which is sometimes explained by evolutionary medicine by reduced selective pressure against cancer in the post-reproductive part of the population. However, certain types of cancer such as testicular cancer with a peak onset between 25 and 34 years ^374,375^ contradict these ideas and there is no consensus yet on this topic. Moreover, there is a steep decline of cancer incidence in very old people, e.g. there is hardly any colon cancer above 80 years of age ^376^. Interestingly, in projects such as the “Zincage study” and others ^377 378^ it has been observed that dietary zinc intake and plasma zinc concentrations decline in the elderly population or see Table 1 in ^379^, which may create a selective force as putative prerequisite for early cancer development (Figure 2). It seems worthwhile to systematically test whether there is a causal link between zinc status and cancer incidence in the aging population.

Naked mole rats are used as a model in aging research and an apparent absence of tumors has been observed in this rodent ^380^. Recently it has been suggested that this phenomenon may be related to their elevated tissue concentrations of high-molecular-mass hyaluronan ^381^. Interestingly, hyaluronic acid forms complexes and interacts with different metal ions such as zinc and copper ^382–384^. Whether there is a link between this characteristic of hyaluronic acid and cancer in naked mole rats remains to be tested, but it is tempting to speculate that the high concentrations of hyaluronic acid may form a protective layer that buffers trace element concentrations in such a way that tumors do not develop.

As for two neurological disease of the elderly population, an inverse correlation in disease risk with different types of cancer has been observed, namely for Alzheimer ^385^ and Parkinson ^386^ disease. Unfortunately, the link has remained enigmatic. Nevertheless, it has also been observed that both diseases show complex patterns of altered trace element levels in different regions in the brain ^387^ and it has been suggested that perturbations in trace element homeostasis may contribute to cognitive loss and neurodegeneration ^388^. Moreover, a causal effect of serum iron levels on Parkinson disease risk has been suggested ^389^. Combined these observations with the AMTC hypothesis, we speculate that deviations from optimal trace element levels in opposite directions may help explain the inverse correlation between cancer and these devastating neurological diseases in the future. In any case, much work remains to be done in order to understand the complexities of trace element homeostasis in the brain and the directions of change that may modify the course of these diseases.

### Additional risk factors in cancer

A. Alcohol consumption and smoking are important independent risk factors for esophageal cancer and they confer further increased cancer risk in combination ^390^. It is still not entirely clear why alcohol increases the risk for a number of cancer types although it was suggested that reactive oxygen species and possibly its effect on other nutrients or its role in the immune system may play a role ^391,392^. In our model of carcinogenesis, smoking would act as the carcinogen and hereby would increase the number of mutated cells upon which selection can act on, while alcohol would trigger the selective force through its potential ability to exacerbate or induce zinc deficiency. The effect of ethanol on zinc levels has been shown in rats ^393–395^ and humans ^396^. An interaction between vitamin A and zinc may play an important modifying role here ^397–399^. Compatible with this picture is the observation, that only the combination of ethanol and a carcinogen (but not alone) on the background of zinc deficiency was sufficient to induce esophageal tumors in rats ^400^. Fittingly, it was suggested that aneuploidy predisposes to certain cancers ^401^. Ex-smokers have only a slightly elevated risk for lung cancer ^402^, an observation which may possibly be explained by purifying selection leading to decreased mutation burden in lung tissue in the absence of carcinogens from cigarette smoke.
B. Cancer incidence and survival shows marked differences between different ethnic groups ^403^. As an example, African Americans show an increased incidence of numerous types of cancer ^404^. A number of known factors such as lifestyle, diet, socio-economic status and access to diagnostic and therapeutic interventions play a role here. Considering the AMTC hypothesis, it may be fruitful to ask whether potential ethnic differences in trace element homeostasis exist and whether they may help explain some of the observed differences. In this direction, differences in serum levels of zinc and iron between African American and Hispanic children have been reported ^405^. Also, a functional amino acid polymorphism (L372V) in the intestinal zinc uptake transporter ZIP4 that shows extreme frequency differences between African and Eurasian populations has been described ^406^, but overall the data are very limited. Similar questions regarding trace element homeostasis could be asked related to the cancer incidence discrepancies that exist between men and women ^407^.
C. Women working night shifts over many years have an increased incidence of endometrial and colorectal cancer^408,409^. Intriguingly, zinc seems to be regulated in a circadian manner in humans ^410^ and rats ^411^. Likewise, serum iron is strongly influenced by the circadian clock and shifted by day sleep patterns ^412^. Whether the perturbance of normal trace element homeostasis in night shift workers may predispose to certain types of cancer could be tested.
D. Observations from the field of medical geology ^413^ have indicated a higher incidence of certain cancer types in trace element deficient regions in China ^414^ and Turkey ^415^ among other countries. While there are many complex interactions between the trace elements along the food chain and while the dietary uptake of elements is currently poorly understood, a causal link between geologically influenced trace element levels in the human diet and cancer incidence should not be ruled out.
E. A number of human diseases lead to an altered cancer incidence. Hereditary hemochromatosis, a disease marked by iron overload in different tissues is linked to an increased risk of liver cancer and other malignancies ^416–418^. Wilson’s disease leads to copper overload in liver through the malfunction of copper transporter ATP7B and different therapies exist that act to normalize copper homeostasis. Cases of hepatocellular carcinoma in Wilson’s disease are known, but evidence of an increase or decrease of cancer risk in this disease seems inconclusive. People with Down syndrome have a sharply increased risk for leukemia while it is reduced for most solid tumors ^419,420^. Interestingly, iron and zinc concentrations in different fractions of blood differ in people with Down syndrome and controls ^421,422^. Colon cancer risk is strongly increased in patients of *Colitis ulcerosa* ^423,424^ and compatible with an early selective force related to tissue zinc deficiency, zinc concentrations were reduced in mucosal biopsies from patients with active inflammatory disease ^425^. Perturbations of trace elements in blood serum are also commonly seen in inflammatory bowel diseases ^230,426^. In summary, there exist a large number of highly diverse risk factors in cancer for which we describe established or suggestive links to trace element homeostasis. This strongly calls for follow-up studies to further investigate the role of trace elements for these cancer risk factors.

## Supplemental Notes

### Supplemental Note A: Material and Methods

#### Data

The fifteen used datasets ^100,117,427–440^ (Supplemental Table S1) consisted mainly of epithelial carcinomas including a collection of cell lines ^437^ which mainly but not exclusively were derived from epithelial cancer specimen. The used datasets are not embargoed according to the TCGA publication guidelines (http://cancergenome.nih.gov/publications/publicationguidelines). For the majority of available copy number data in cBioPortal, the GISTIC 2.0 algorithm ^30^ and in some cases the RAE algorithm ^427^ have been used. This copy number information should be considered putative due to factors outside the reach of the authors, namely technical challenges in calling correct copy numbers. For example it is difficult to differentiate between gains and amplifications due tumor heterogeneity and tumor impurity ^427^. Gene expression levels were inferred through microarray or RNA sequencing technology as indicated in Supplemental Table S1. Current datasets such as those from the Cancer Genome Atlas Project have limitations with regard to the number of analyzed cases and a bias towards late stage tumors while analyses of early or pre-cancerous specimen are largely missing. Nevertheless, we think that the datasets are sufficiently accurate for our analyses and conclusions. The analyzed genes (N=458) included zinc, iron and copper transporter genes and additional, manually curated genes that are related to trace elements; all remaining solute carrier (SLC) transporter genes which were used as controls (N=326); a group of established cancer genes (N=52) from the Cancer Gene Census ^107^, that included all genes denoted as large deletions and amplifications (accessed in June 2013) plus six additional cancer genes ^19^ (N=6; *BRAF*, *CDKN2B*, *MET*, *SMAD2*, *TERT*, *TP53*). Data equivalent to the data available at cBioPortal for Cancer Genomics ^105,106^ was downloaded on July 7^th^, 2013, using the Cancer Genomics Data Server (CGDS), hosted by the Computational Biology Center at Memorial-Sloan-Kettering Cancer Center (MSKCC) and using the CGDS-R library by Anders Jacobsen.

In order to obtain information on point mutations as well as additional gene information, all genes were queried through the intOgen ^108^ TCGA portal with the options “Drivers - View matrix of candidate drivers per cancer type” using PANCANCER12 as cancer type. Interestingly, a recent study ^84^ found that many of the currently suggested candidate driver genes may represent spurious findings that were confounded by marked heterogeneity of mutation rates (due to correlations with low expression and late replication time) along the genome. intOgen results are largely robust to the heterogeneity of mutation rates due to their “within gene” tests of positive selection. Results on point mutations were enriched with results from MutSigCV ^84^ and MuSiC ^441^. Additional information on diverse mutations including amplifications and deletions was included from Cancer Gene Census ^107^ and shown in Supplemental Table S2. Focally amplified and deleted genes based on TCGA data were downloaded from the Broad Institute TCGA Genome Data Analysis Center (Broad Institute of MIT and Harvard: doi:10.7908/C1736NW2; analysis from April 21^st^, 2013). In the copy number analysis (GISTIC2) based on the PANCANCER cohort with 12 disease types and 4976 tumor samples, the identified 41 significant focal amplifications and 51 significant focal deletions contained 10 and 12 genes in our set of 58 prioritized cancer genes, respectively. Information from individual cancer types was not included, which may have further increased the overlap between analyzed cancer genes and the focally aberrated regions.

#### Analysis

A correlation between mRNA expression and copy number was calculated for each pair of gene and cancer type. Genes in a specific cancer type with a Pearson correlation > 0.1 between mRNA expression and copy number resulting in a p-value < 0.05 were considered to show a gene dosage effect (recorded in a binary fashion, TRUE or FALSE). Only samples with one, two or three copies of a certain gene were considered. Implicit in the Pearson correlation coefficient, we assume a linear correlation between gene copy number and expression although gene dosage in many cases is more complex ^442^. Percentage of genes with gene dosage (Supplemental Figure S3, Supplemental Tables S4 and S5) was calculated for each dataset separately as the number of genes with gene dosage divided by all analyzed genes.

For each pair of gene and cancer type, a score was calculated that builds on three pieces of information considered as meaningful criteria for candidate driver genes through aneuploidy:

score_1=(maximal frequency of gains or losses) * [log (gains/losses)] * (gene dosage)

Samples showing zero gains or losses were set to one in order to avoid infinite values. These scores were made comparable across cancer types by dividing through the mean score_1 of all SLC control genes for a particular cancer type, e.g.:

score_1_corrected_Melanoma=score_1 / mean (score_1_Melanoma of SLC genes 1 to 326)

The mean of this score was calculated across different groups of genes for each cancer type. The resulting data points from twelve epithelial cancers were used for the comparison of zinc transporters and established cancer genes against the group of SLC control genes (Figure 3; Supplemental Table S6).

For each gene, different aneuploidy scores were calculated:

aneuploidy_score_1 = median (score_1_corrected_cancer_type_1, score_1_corrected_cancer_type_2, …, score_1_corrected_cancer_type_12)
aneuploidy_score_2 = aneuploidy_score_1 * square_root (number of counted dosage effects out of 12 epithelial cancers)

This score seems useful to compare genes with other genes and was used in Tables 1A and 1B, assuming that strong candidate driver genes for epithelial cancers show gene dosage across many different types of epithelial cancer.

aneuploidy_score_3=(mean(maximal_frequency_of_gains_or_losses in epithelial cancers with gene dosage)) * (mean([log (gains/losses)] in epithelial cancers with gene dosage)) * square_root (number of counted dosage effects out of 12 epithelial cancers)

This score is an alternative, possibly less powerful aneuploidy score which was not further used in this manuscript. In Supplemental Table S2, “hetloss” stands for 1 gene copy (hemizygous loss), "e_p" stands for "epithelial cancer, published" and "o_p" stands for "other type of publisehd cancer type". A provisional R script is available on request.

The median frequencies of gains or losses for each gene across epithelial cancer types are listed as “aberration frequency across epithelial cancers - only gene dosage”. The median ratios of gene losses versus gains in epithelial cancer types (again only using correlations with gene dosage) were used as a proxy for the tendency of a gene to be gained (median ratio >2), ambiguous (2>median ratio>0.5) or lost (median ratio<0.5).

In the future, this analysis could certainly be improved by correcting for chromosome arm sizes or sample sizes, by including a more comprehensive set of established cancer genes or by an unbiased genome-wide approach as well as by including additional datasets and different layers of functional data.

#### Additional notes on the analysis

It is interesting to note that a number of zinc-related genes such as some metallothionein genes on chromosome arm 16q12-13 or iron-related genes such as *RHOA*, *FECH*, *BDH2*, *LTF* or *HIF1A* or copper-related genes such as *CP*, *ATOX1*, *COX17* or *COMMD1*, the magnesium transporter gene *SLC41A1* and cation channel gene *TRPM7* as well as numerous SLC transporter genes from various families reach high aneuploidy scores as well. Whether some of these genes may play a role as drivers in aneuploidy remains to be tested. However, it is important to note that more likely, many of these genes represent mere “hitchhikers” that cluster in recurrently gained or lost regions such as chromosome arm 1q. Analogous to genetic hitchhiking which confers neighboring alleles of an selected advantageous allele similar genetic signatures ^443^, in cancer clones in the absence of meiotic recombination and in the presence of large, chromosome-arm gains and losses, one can expect complete linkage between selected and hitchhiking alleles. Such effects are likely to blur the differences and only through the observation of convergent aneuploidic events in hundreds or thousands of cancer specimens we may hope to see a small statistical difference between selected and “hitchhiking” genes. The copper transporter *ATP7A* on chromosome X was not accessible in most of the datasets and should be investigated in the future.

*MYC* and *SLC39A4* (ZIP4) are both located on chromosome band 8q24, at a physical distance of around 15 Mb. mRNA expression levels can be plotted against gene copy number (http://www.cbioportal.org). *MYC* seems to show a stronger gene dosage effect due to high level amplifications, whereas the increase of *SLC39A4* mRNA expression is mostly by chromosomal gains. Since the overexpression of SLC39A4 (ZIP4) may contribute to oncogenesis ^56,58,189,444^ as was shown for MYC ^445–447^, both may act as contributing or even driving factors.

It was described that almost all zinc transporters are downregulated in pancreatic cancer on the mRNA level, except for *SLC30A4* ^448^. A similar pattern was found for selected zinc transporters in prostate cancer ^449^. This may likely be a consequence of overall reduced zinc levels in the cancerous tissue. This deregulation of zinc transporters may also conceal any possible link between mRNA or protein expression of zinc transporters between healthy and malignant tissue. These observations do not contradict our model, since gene dosage effects are still observable between samples of varying copy number in tumor samples.

### Supplemental Note B: Limitations and complexities of the AMTC hypothesis

First, the AMTC hypothesis is based on diverse observations from zinc biology and a tentative analysis of genomic data derived from a limited number of tumor samples, and is currently not further supported by direct experimental or epidemiological evidence. It should thus be regarded merely as theoretical working model that remains to be tested in a rigorous manner.

Second, alternative explanations for the observed correlation between trace element deregulation and chromosomal gains and losses in cancer might exist. While it seems clear that trace elements indeed play important roles during carcinogenesis, their deregulation does not necessarily have to be causal and the link to chromosomal gains and losses that we propose remains largely untested. Moreover, in light of trace elements being involved in many different cellular processes, the correlation between deregulated trace element homeostasis and cancer development could be coincidental rather than causal.

Third, other candidate genes, including established tumor suppressor genes and oncogenes, have recently been suggested as potential drivers of the observed patterns of chromosomal aberrations^111^. While we by no means wish to dispute their importance during carcinogenesis, we note, however, that most of these cancer genes are frequently affected by focal mutations or point mutations and that therefore their causal contribution to the chromosome scale events leading to aneuploidy remains to be explored.

Fourth, not all cancer types seem to accumulate chromosomal gains and losses at high frequencies, e.g. different types of leukemia and lymphoma. In breast cancer, the subtype "IntClust 4" was characterized as a ‘CNA-devoid’ subgroup with frequent lymphocytic infiltration ^110^. This may hint to a prominent role of other mechanisms such as microRNAs ^450^ or the effect of point mutations. With regard to the AMTC hypothesis, it is unclear whether trace elements might also play a role in these types of cancer.

Fifth, if metal transporters are indeed potential drivers of aneuploidy, it may seem surprising that metal transporter genes are not frequently enriched in point mutation (e.g. loss of function) or focal aberration analyses in recent resequencing projects. At the same time, it seems possible that gene expression changes through non-focal chromosome-arm losses or through focal events in regulatory regions would be more important, and such regulatory regions currently are poorly annotated and thus largely missed by resequencing approaches.

Sixth, even if this hypothesis turns out to be valid, the directions of the resulting changes in intracellular and intra-organellar zinc concentrations due to chromosomal gains or losses would remain complex. Moreover, it would not be immediately clear if one should expect reduced cellular zinc influx and increased zinc efflux - which may explain the observed reduced zinc levels in most cancerous tumors - or rather the opposite, e.g. increased zinc influx that may confer a fitness advantage to the pre-cancerous cell under a possible local zinc deficiency in the tissue. This is complicated by the fact that for most metals, there are influx and efflux transporters that target different organs and cell compartments. Further, metal transporters show different degrees of substrate affinity. As an additional complication, there are different zinc pools (zinc and zinc bound by different proteins). For copper there are also labile pools and pools of copper bound by metallothionein or ceruloplasmin. In this context it should be pointed out that zinc is not always deregulated in the same direction in cancer tissue. While the overall chromosomal aberration pattern is similar to other epithelial cancers, breast cancer tissue shows higher concentrations of zinc (opposite to prostate and pancreatic cancer tissues). However, this may be reconciled by a closer look at the different existing zinc pools in breast cells, and it might be possible that while the tissue zinc concentration is increased, a different zinc pool such as possibly the intracellular pool of labile zinc, indeed follows the same direction as other tumors do.

Seventh, even if correct, it is almost certain that this picture is still incomplete. For example, it seems likely that there are many unknown regulators of trace element homeostasis still to be found, or that known regulators were missed in our analysis. This may include long non-coding RNA (lncRNA), microRNAs mir182 ^451^ or miR-31 and miR-21 ^452,453^, the zinc sensing receptor GPR39 ^454^ and unknown regulators. Other important players in zinc homeostasis are the interleukin IL-6, which interacts with ZIP4 ^332^, KLF4 as regulator of ZIP4 ^194,195^ and the copper zinc superoxide dismutase (SOD1) ^455^. Other ion channels may be more unspecific and possibly have a role in zinc homeostasis, such as TRPM7 ^456^ or MCOLN1 (TRPML1) ^457,458^. Promoter methylation of the metallothionein MT1G was suggested as a marker for tumor aggressiveness in prostate cancer ^459^. In summary, additional genes likely play a role in the regulation of trace element homeostasis and possibly in cancer related aneuploidy. In this context we note that we are unable to present strong candidate drivers for some of the recurrent aberrations, for example of chromosome 20.

Eighth, the identity of a possible driver gene behind the recurrent gain of the long arm of chromosome one (+1q) seems somewhat unclear. Several ion transporter genes are located on 1q, including zinc efflux transporters *SLC30A1*, *SLC30A10*, magnesium transporter *SLC41A1* and zinc influx transporter *SLC39A1*. Although *SLC39A1* reaches the highest aneuploidy scores and shows gene dosage in the highest number of epithelial datasets (Table 1), experimental results possibly conflict with a role of *SLC39A1* gains in carcinogenesis, as overexpression inhibits tumor growth ^64,65^. Hence, other genes including *SLC30A1* or *SLC30A10* might be better candidate drivers in all or selected cancer types. Nevertheless, it is also worth pointing out that SLC39A1 is an important zinc transporter shown by its near-ubiquitous expression pattern ^460,461^, which again would favor SLC39A1. The property of SLC39A1 as to promoting zinc uptake might render it a candidate as very early chromosomal gain mutation and possibly allow the pre-cancerous cell to overcome growth control or reach a fitness advantage in a hypothesized local zinc deficient tissue. In this direction, in a study on breast cancer, the sole presence of +1q indicated that this was an early event. Also in breast cancer, in a recent sophisticated analysis the temporal order of diverse mutation events was disentangled and gain of 1q was resolved as first mutational event in some cancer specimen ^462^.

Ninth, although in this article we discuss arm-level and focal mutations separately, they may not be entirely independent. For example, focally amplified regions seem to occur on gained chromosome arms but never on hemizygously lost chromosome arms and *vice versa* with focal deletions, as observed visually in Figure 1 in ^438^ and Figure 2 in ^463^. Does this correlation mean that one-copy gains and multi-copy amplifications are from the same mutation spectrum and caused by the same underlying driver genes? Fittingly, in many CGH studies, a “minimal region of overlap” between genetic markers was defined in order to identify the smallest possible chromosomal region that likely harbored the suspected driver gene. However, this interpretation may have blurred the possibly meaningful distinction between drivers of arm-level versus focal events. Alternatively, may this correlation mean that one type of mutation facilitates the emergence of the other? If so, what type of mutation comes first? The temporal order of mutations is difficult to resolve, but some results exist. For example, focal events have been observed to occur late as compared to arm-level events in a study of early versus advanced ovarian cancer ^464^.

Tenth, we ask if such a simple hypothesis may have possibly been overlooked? On the one hand, the answer may be “yes”, since no attempts have been made so far to jointly integrate trace element biology and evolutionary genomics in cancer research. Trace elements are understudied, no strong biomarkers for human zinc status exist and ionomic approaches are not well integrated with the other omic technologies in humans. For example, only few genome-wide association studies have been carried out for the trace elements iron, magnesium, copper, selenium and zinc in human serum or erythrocytes ^465–469^ and none so far in human tissue. Moreover, this idea may have evaded discovery because of the functional redundancy of many metal ion transporters. Despite all efforts, “cancer genomics is in its infancy” ^470^ and possibly, while recent efforts in cancer genomics have successfully focused on point mutations and focal amplifications and deletions, the underlying causes for the observed aneuploidy patterns have remained difficult to determine.

### Supplemental Note C: Trace element homeostasis and cancer

In living organisms, essential trace elements are tightly regulated. Their concentrations are kept on a narrow plateau; both their deficiency and excess are detrimental ^471^. A potential role of trace elements in cancer has been observed early ^472^, and has been blamed on DNA damage induced by micronutrient deficiencies by some authors ^473^. The status of essentiality for humans or other organisms has only been recognized in a long process and for each element individually ^471^. Some essential trace elements commonly show deficiency symptoms in humans. For example, symptoms of zinc deficiency in humans were first discovered and described by Ananda S. Prasad in the 1960s ^474^.

Some important metal transporters are the following. With regard to zinc, the *SLC39A* gene family encoding Zip1-Zip14 and the *SLC30A* gene family encoding ZnT1-ZnT-10 control the cytosolic and organellar levels of zinc ^475^. SLC30A10 is a manganese transporter ^476,477^. Copper is controlled by four different transporters (SLC31A1, SLC31A2, ATP7A and ATP7B). Iron homeostasis is complex, but the number of human iron transporters (TFRC, TFR2, SLC11A1, SLC11A2, SLC40A1) is similarly low as in copper. Different metal-binding proteins such as the metallothioneins (zinc and copper), ceruloplasmin (copper), albumin (zinc and copper), hemoglobin (iron), transferrin (iron), lactoferrin (iron), a number of microRNAs, the metal regulatory transcription factor MTF-1 and a number of known and unknown additional regulators contribute to maintaining the homeostasis of trace elements. This complex regulation extends over different time scales and includes all cellular compartments as well as diverse types of cells, tissues and organs. There are numerous metal-binding proteins in humans including metalloenzymes and structural proteins. Estimates for zinc-binding proteins reach up to 2800 ^478^, many of which are transcription factors including zinc-finger proteins. There are manifold interactions between different trace elements and some ions are transported by the same transporters, e.g. SLC39A14 which transports both zinc and iron ^479^. Specifically, the antagonism between zinc and copper ^480,481^ could mean that either change (zinc or copper) may disrupt the concentrations of the other. This was demonstrated recently in mouse model of the zinc-deficienct disease acrodermatitis enteropathica, in which the induced organismal zinc deficiency on the long run led to an accumulation of copper, iron and manganese in liver ^482^.

Zinc has been extensively discussed in the main manuscript, so here we just point out a few additional results from the literature. For example, SLC39A4 (ZIP4) was suggested as a marker for survival in Glioma ^483^. Heat shock proteins are highly expressed in cancerous cells, which contributes to their survival ^484^, and interestingly, they are also upregulated zinc or zinc-pyrithione treated epidermal cells ^485^. In a genome-wide study, the mRNA expression pattern of *SLC30A4* was found to stand out and the corresponding protein ZnT4 was found to decrease from benign to invasive and to metastatic prostate cancer ^486^. Zinc transporters in prostate cancer have been reviewed recently^487^. In genome-wide association studies a regulatory variant (rs1050631) was identified that modulates the expression of zinc transporter SLC39A6 and which was associated to survival in esophageal cancer ^488^.

The non-essential, toxic trace element cadmium can be found just below zinc in the periodic table and behaves similarly in many biochemical reactions. Cadmium acts as an antagonist of zinc, has received a lot of attention and also has been suggested to be a cause of cancer ^489^. For example, toe nail cadmium was associated with an increase in prostate cancer risk ^490^. While cadmium has no physiological role and is toxic at the best, its antagonist role of zinc may actually explain some of its cancerogenic potential. As was pointed out ^489^, “the substitution of cadmium for zinc may be a central mechanism underlying the carcinogenicity of cadmium”.

As for copper and cancer, much fewer studies have been carried out. Importantly and as mentioned in the main text, copper concentrations are often increased in cancerous tumors. Of note, platinum-based anti-cancer drugs use the cellular copper transport. An overview on cisplatin resistance and these three copper transporters is given here ^491^.

Different links between iron homeostasis and cancer have been described in an early ^492^ and recent review ^493^. On link relates to hemochromatosis, a genetic disease characterized by iron overload, especially in the liver. Interestingly, not only iron but also zinc and possibly other trace elements seem to be deregulated ^494^. Hemochromatosis patients show an increased risk for diverse cancers such as liver cancer ^495^, gastric cancer ^496^, breast and colorectal cancers ^497^. While the mechanism for the association between the hemochromatosis mutation HFE C282Y and cancer is not entirely clear, it is most likely related to iron ^498^. Due to its association with hemochromatosis, iron was described as a “carcinogenic” metal ^499^.

There is a strongly debated link between selenium and different types of cancer. Patients of lung and laryngeal cancers show a significant decrease of serum selenium ^500^. Selenium supplementation was seen as a promising preventive approach against different cancers, although a large recent clinical trial failed to find an effect ^501^. However, this was possibly due to the fact that the American cohort was selenium-sufficient and not selenium-deficient and other reasons. Selenium is incorporated by selenoproteins in form of selenocysteine and 25 selenoproteins have been found in humans ^502^. SEPP1 contains 16 selenocysteins, is produced by the liver and is a main player in the distribution of selenium within the body. SEPP1 ^503^ and several other selenoproteins such as SEP15 ^504^ have been implicted in cancer.

### Supplemental Note D: Anecdotal evidence from comparative oncogenomics

As suggested in Text Box B, comparative oncogenomics could be used to test the AMTC hypothesis for compatibility with the existing data. The underlying assumption is, that genes possess similar functions in different organisms and that therefore, the observed aneuploidy patterns for a particular type of cancer are driven by the same driver genes. Here, we limit ourselves to a few superficial, almost anecdotal observations. As an example, a study based on array CGH compares gain and loss patterns between dogs and humans for osteosarcoma highlights four overlapping regions corresponding to human 1p36, 1q21-22, 6p22 and 8q24 (Figure 5) ^505^ which possibly might imply *SLC39A1* and *SLC39A4* (or *MYC*). In osteosarcoma from a third species, mouse ^506^, 3696 identified overlapping genes include the metal transporters *SLC30A8*, *TFR2*, *SLC39A1* and *SLC39A4*. Hence, *SLC39A1* and *SLC39A4* (and *MYC*) seem to be gained in osteosarcoma in all three species, human, mouse and dog. In a different comparison with zebrafish malignant peripheral nerve sheath tumors, Supplemental Dataset S5 from ^507^: of 866 shared gained genes (“all-all” comparison), *SLC39A4* and *SLC30A8* represent metal transporters, and of 1774 shared lost genes, *SLC39A14* and *SLC39A6* represent metal transporters. *MYC* was not identified as a shared gene. There are many more studies that could be used but this would be out of scope for this article.

### Supplemental Figures

**Supplemental Figure S1:**
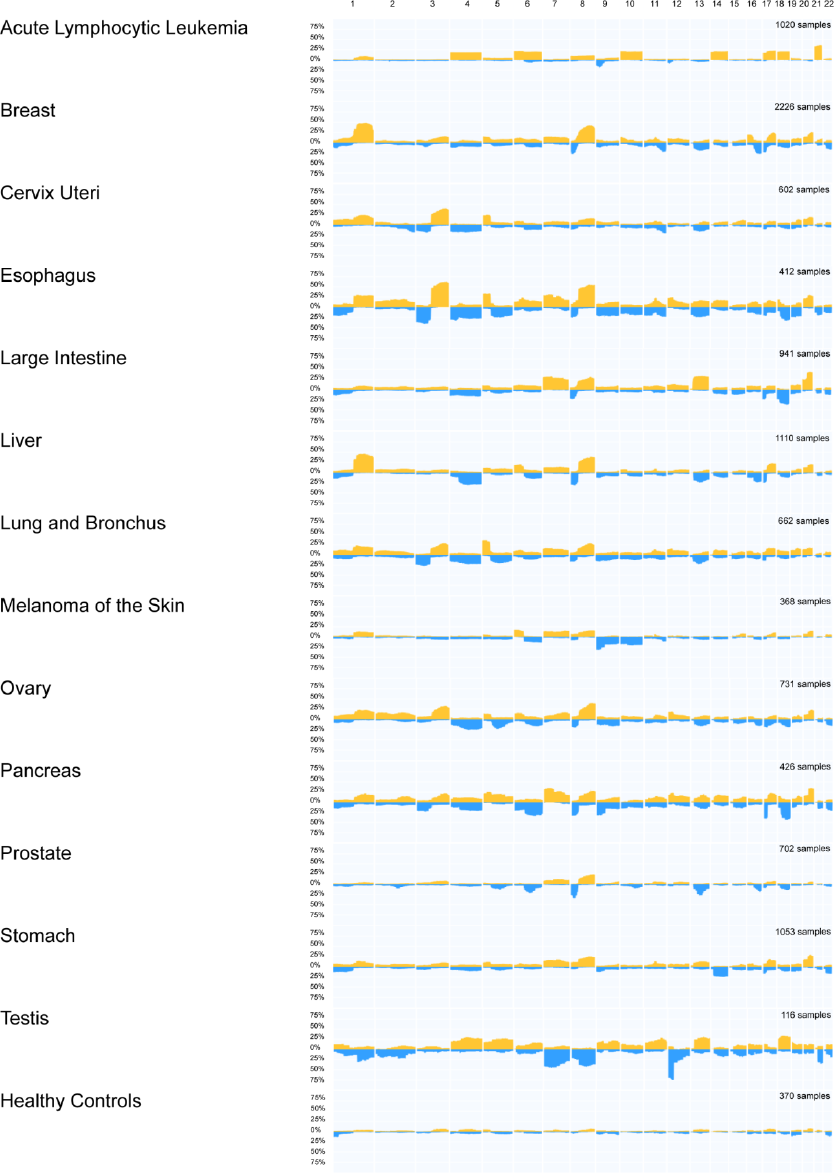
Additional aneuploidy profiles of different cancer types using the SEER nomenclature. Percentages of gains (orange) and losses (blue) across analyzed samples are shown on a scale from 0%, 25%, 50%, 75% to 100%. Note the near-absence of aneuploidy in healthy controls. Plots were generated at http://progenetix.org.

**Supplemental Figure S2A:**
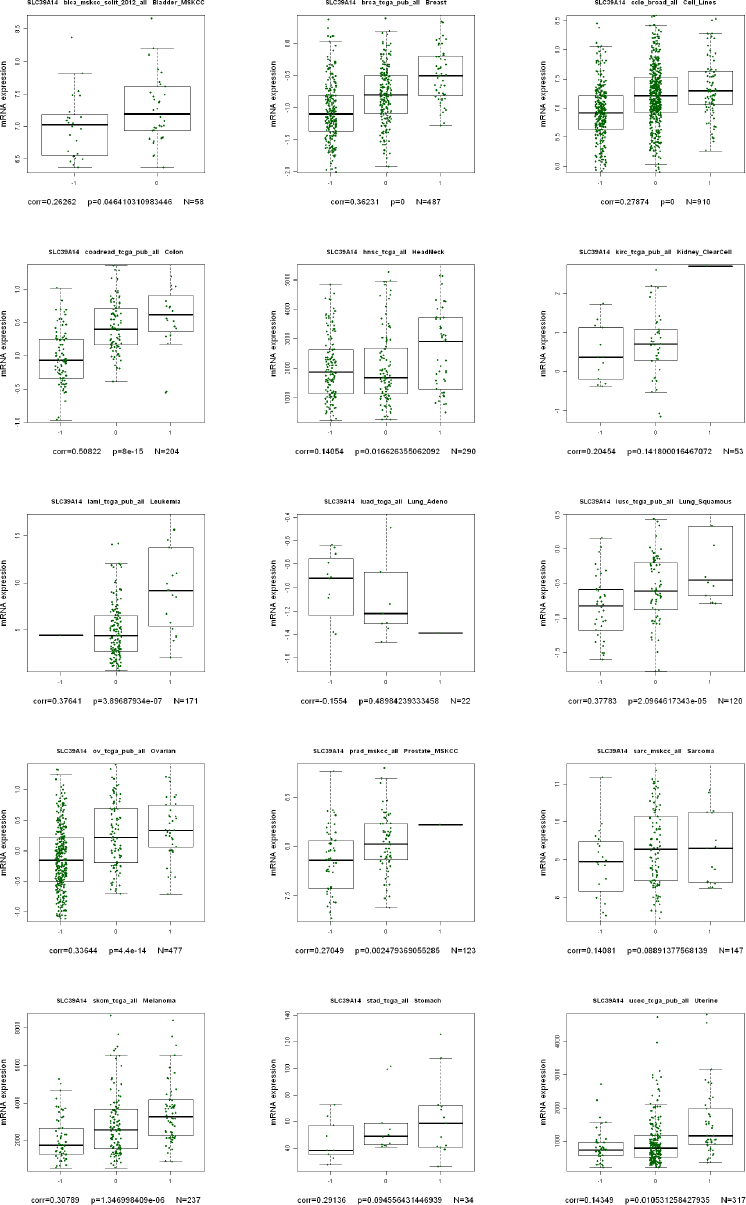
Gene dosage analyzed in all fifteen datasets; zinc transporter SLC39A14.

**Supplemental Figure S2B:**
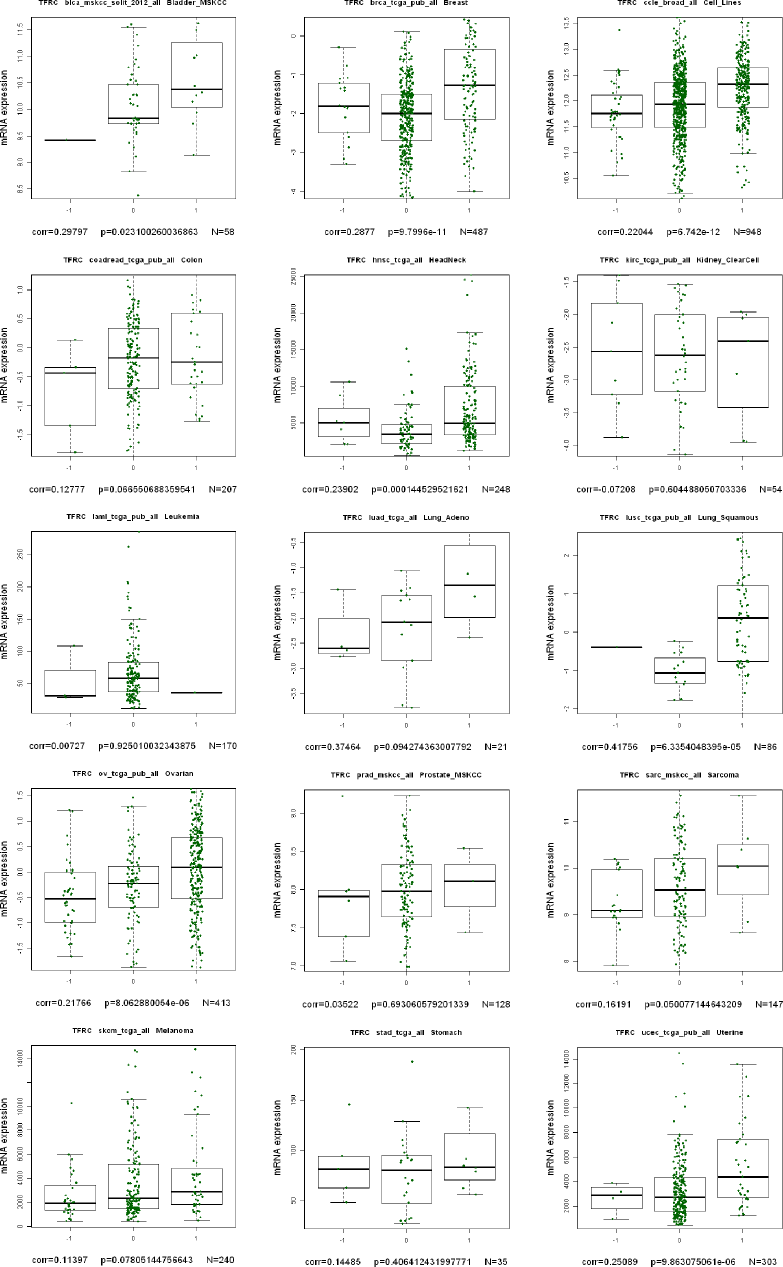
Gene dosage analyzed in all fifteen datasets; iron transporter TFRC.

**Supplemental Figure S3:**
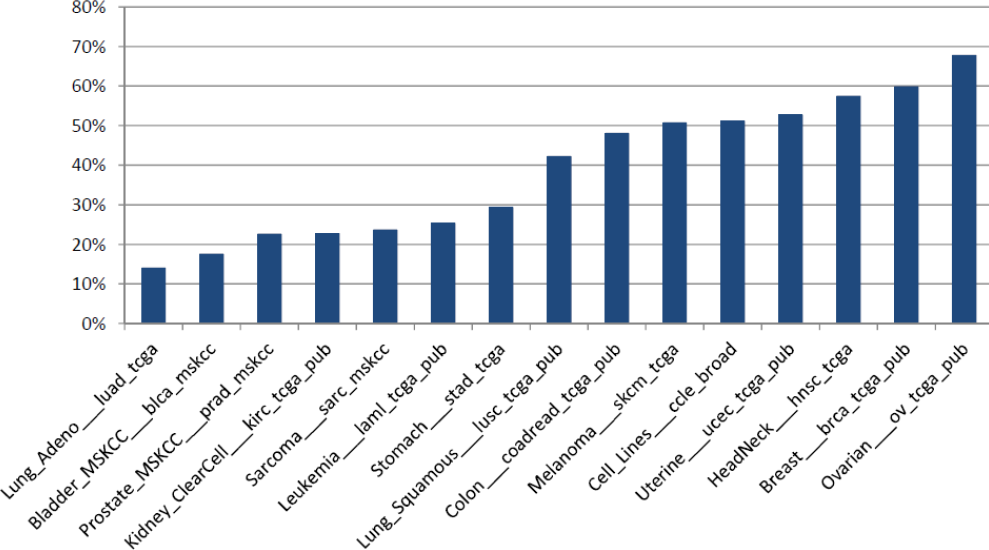
Percentage of genes with significant gene expression dosage relative to all 458 analyzed genes in the 15 analyzed datasets. The assignment of gene dosage is sensitive to parameter choice and the number of samples analyzed within a study.

